# Contribution of amygdala to dynamic model arbitration under uncertainty

**DOI:** 10.1101/2024.09.13.612869

**Authors:** Jae Hyung Woo, Vincent D. Costa, Craig A. Taswell, Kathryn M. Rothenhoefer, Bruno B. Averbeck, Alireza Soltani

## Abstract

Intrinsic uncertainty in the reward environment requires the brain to run multiple models simultaneously to predict outcomes based on preceding cues or actions, commonly referred to as stimulus- and action-based learning. Ultimately, the brain also must adopt appropriate choice behavior using reliability of these models. Here, we combined multiple experimental and computational approaches to quantify concurrent learning in monkeys performing tasks with different levels of uncertainty about the model of the environment. By comparing behavior in control monkeys and monkeys with bilateral lesions to the amygdala or ventral striatum, we found evidence for dynamic, competitive interaction between stimulus-based and action-based learning, and for a distinct role of the amygdala. Specifically, we demonstrate that the amygdala adjusts the initial balance between the two learning systems, thereby altering the interaction between arbitration and learning that shapes the time course of both learning and choice behaviors. This novel role of the amygdala can account for existing contradictory observations and provides testable predictions for future studies into circuit-level mechanisms of flexible learning and choice under uncertainty.

## Introduction

One of the challenging aspects of learning in naturalistic settings is that it is inherently unclear what features or attributes of a choice option are predictive of subsequent reward outcome. Imagine successfully operating a new coffee machine after switching it on and pressing a flashing button on the left side of its screen. What should you press the next time you want to get coffee: the same button or any button that is flashing? In this specific example, reward outcomes can be equally attributed to either the identity of a choice option (e.g., flashing button) or the action needed to obtain that option (pressing the left button), corresponding to uncertainty about the correct model of the environment (stimulus-based vs. action-based). More generally and during most naturalistic settings, reward outcomes could be linked to any features and/or attributes of a selected option or chosen action. It has been suggested that the brain tackles such uncertainty by running multiple internal models of the environment, each predicting outcomes based on different attributes of choice options, and using the reliability of these predictions to select the appropriate model to inform choice behavior^1–3^.

Although there exist many conceptual and algorithmic solutions to model arbitration^2–4^, confirming implementation level details in terms of the operation of neural circuits has remained a challenge due to several factors. First, most experimental paradigms involve manipulating uncertainty in one of two ways. In some paradigms, there is uncertainty about which of multiple choice options is more rewarding through probabilistic reward contingencies and contingency reversals^5–7^. In other paradigms, the more rewarding option is easily ascertained, but there is uncertainty about the correct model of the environment and when to choose that option^8–10^. Few if any studies have manipulated uncertainty about stimulus-action-outcome relationships in conjunction with uncertainty about which model of the reward environment is currently relevant. In reward environments where the correct model of the environment does not change frequently, reliabilities of different models can reach their asymptotes very quickly, concealing the contributions of circuits involved in dynamic arbitration and model selection. Second, it is intrinsically difficult to measure and track the contributions of multiple learning systems and their interactions because different learning systems can drive choice behavior at any given moment. Third, because computations required for learning and arbitration under uncertainty must interact with each other, many cortical and subcortical areas may appear to be similarly involved in different processes (lack of specialization). For example, the amygdala has been shown to contribute to reward learning under uncertainty by both improving and impairing learning performance^11–17^ and by its contribution being associated with different types of uncertainty^18,19^.

To overcome these challenges and reveal the circuit and neural mechanisms underlying arbitration, we applied multiple experimental and computational methods to examine choice behavior in three groups of monkeys (control group and two groups with bilateral lesions to the amygdala or ventral striatum) performing a probabilistic learning task that involved multiple forms of uncertainty. These included uncertainty about the better option, uncertainty about the correct model of the environment, and uncertainty about when reward associations change, thus creating a challenging task that could reveal the role of the amygdala and ventral striatum in all three of these processes. To track the simultaneous contributions of multiple learning systems and their interaction, we extended metrics based on information theory^20,21^ to quantify consistency in choice and learning based on stimulus- and action-based learning over time. Moreover, we constructed multiple models based on reinforcement learning (RL) to fit choice behavior on a trial-by-trial basis. We examined the best models and their estimated parameters to pinpoint the roles of the amygdala and ventral striatum in reward learning and arbitration. By applying the aforementioned methods, we provide evidence for interactions between stimulus-based and action-based learning, reveal possible mechanisms underlying arbitration between the two systems and how arbitration and learning processes interact, and determine the contributions of the amygdala to this arbitration and the overall learning behavior.

## Results

### Behavioral paradigm with multiple forms of uncertainty

We examined monkeys’ choice behavior when performing a variant of a probabilistic learning paradigm that involves multiple forms of uncertainty. In this paradigm, during each block of 80 trials, monkeys selected between two novel visual stimuli that were randomly presented on the opposite sides of the screen (**Fig. 1A**). Selection of each option was rewarded with a certain probability (80:20, 70:30, or 60:40), but the probabilities for better and worse options reversed at a random point within the block without any signal to the monkey. Critically, unbeknown to the monkeys, reward probabilities for a particular block were exclusively linked with either stimuli or the location at which they were presented (**Fig. 1B**), creating uncertainty about the correct model of the environment (stimulus-based vs. action-based). In one task, rewards were exclusively based on stimulus-outcome associations (What-only task; **Fig. 1C**). In the second task, rewards in each block of trials were determined by either stimulus-outcome or action-outcome associations (What/Where task; **Fig. 1D**). Together, these components resulted in uncertainties about the better option, the correct model of the environment, and when reward associations reverse.

**Fig. 1.**
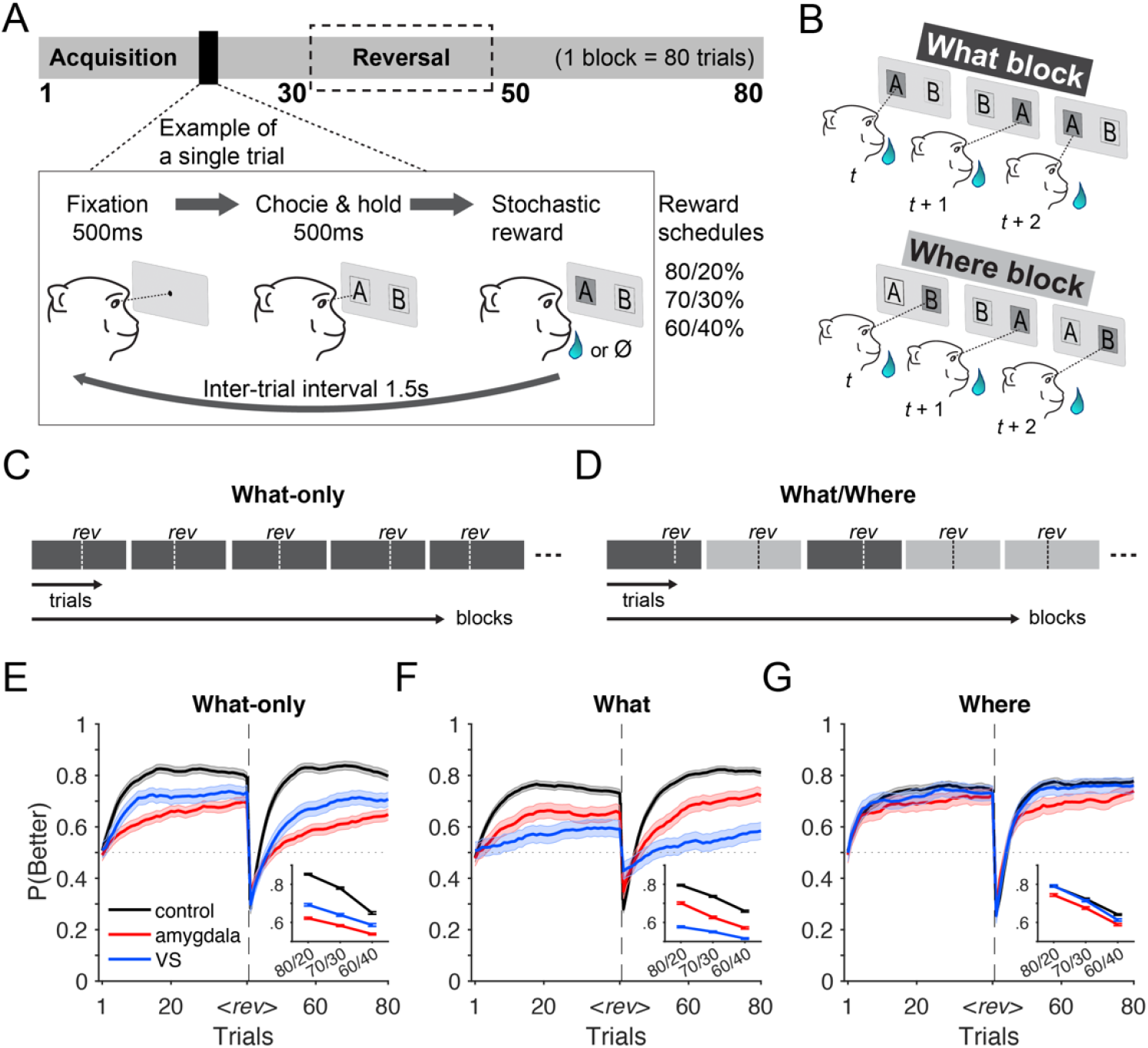
Experimental paradigm, block types, and time course of performance. (**A**) Timeline of each block and a single trial of the experiment. At the beginning of each block, two novel stimuli (abstract visual objects) were introduced. On each trial, animals indicated a choice by making a saccade toward one of the two options on the left and right sides of fixation. Selection of each stimulus was rewarded probabilistically based on three reward schedules. Reward contingencies reversed between better and worse options on a randomly selected trial (between trials 30 and 50). (**B**) Different block types. In *What* blocks (top), reward probabilities were assigned based on stimulus identity, with a particular object having a higher reward probability. In *Where* blocks (bottom), reward probabilities were assigned based on the location of the stimuli, with a particular side having a higher reward probability regardless of the object appearing on that side. (**C**) Outline of the What-only task. Here, only What blocks were used for the entire experiment. Vertical dotted lines, *rev*, indicate a random reversal point within each block. (**D**) Outline of the What/Where task. In this task, two block types, What (black) and Where (gray), were randomly interleaved. (**E–G**) Time course of performance of the monkeys in each group, measured as the probability of choosing the better option, P(Better), separately for different tasks and block types. (Error bars = SEM.) Insets show averaged performance of the blocks by reward schedules.

### Evidence for multiple learning systems and their interaction

To examine the presence of multiple learning systems and reveal the effects of different forms of uncertainty, we first compared the performance of control monkeys across different tasks and reward schedules. Overall performance was best during the What-only task in which only the stimulus identity was predictive of reward, and thus, there was no uncertainty about the model of the environment. The probability of choosing the more rewarding option, P(Better), was significantly higher than that of either What blocks (two-sample t-test; *t*(4674) = 6.415, *p* = 1.54×10^−10^; Cohen’s *d* = 0.196) or Where blocks (*t*(4604) = 9.434, *p* = 6.09×10^−21^; Cohen’s *d* = 0.289) of the What/Where task that involved additional uncertainty about the correct model of the environment. We also observed the effect of expected uncertainty on performance: across all block types and tasks, performance improved as it became easier to discriminate the more rewarding option (*F*(2,7604) = 723.37, *p <* .001).

To capture the effect of reward feedback and how it was used to learn and adjust behavior, we utilized information-theoretic metrics to quantify consistency in reward-dependent choice strategy on two attribute dimensions, stimulus identity and stimulus location^20,21^. Specifically, we examined the conditional entropy of reward-dependent strategy (ERDS), defined as the Shannon entropy of stay/switch strategy conditioned on the previous reward feedback, separately for stay/switch based on action or stimulus identity (see **Methods** for more details). Lower values of ERDS_Stim_ suggest that the animals stayed or switched after reward feedback based on stimulus identity (stimulus-based learning), whereas lower values of ERDS_Action_ indicate that the animals’ stay/switch strategy was based on assigning reward to the chosen action (action-based learning).

To quantify the interaction between the two learning systems, we computed correlation between ERDS_Stim_ and ERDS_Action_ computed from each 80-trials block. We found that even in the What-only task for which action-based learning was irrelevant and minimally used, there was a negative correlation between ERDS_Stim_ and ERDS_Action_ (Spearman’s correlation, *r* = -.196, *p* = 4.89×10^−17^), suggesting that more consistency in using one model resulted in less consistency using the other model. For the What/Where task, we observed stronger negative correlations between ERDS_Stim_ and action ERDS_Action_ for both block types (What: *r* = -.602, *p* = 2.43×10^−296^; Where: *r* = -.578, *p* = 1.32×10^−261^). Overall, these results reveal significant interactions between the stimulus- and action-based learning systems.

Considering previous findings of the influence of learning strategy on the response time^22,23^, we hypothesized that reaction time (RT) depends on the learning system that dominates the behavior at the time. To test this hypothesis, we categorized trials as either stimulus- or action-dominant by directly comparing ERDS_Stim_ and ERDS_Action_ (see **Methods** for details). Using this approach, we found that during the What/Where task, responses in stimulus-dominant trials were significantly slower than action-dominant trials in both block types (**Supplementary Note 1**). This contrast in stimulus vs. action driven RTs was harder to identify in the What-only task in which choices were dominated by the stimulus-based system. Nonetheless, rare action-dominant trials happened when reward value estimates based on the two systems were close to each other, resulting in slower and more erroneous responses. Overall, these results show that entropy-based metrics could be used to identify the adopted model on a given trial and that RT reflected the adopted strategy, with stimulus-based strategy resulting in longer RT than action-based strategy.

### Influences of amygdala and ventral striatum lesions on learning and choice behavior

Next, we compared the effects of amygdala and ventral striatum (VS) lesions on choice behavior to elucidate their contributions to decision making, learning, and arbitration. During What-only task, amygdala-lesioned monkeys exhibited the largest impairment in performance (P(Better) across all three reward schedules: M±SD = 0.581±0.11; **Fig. 1E**). VS-lesioned monkeys also showed impairment compared to the controls (two-sample t-test: *t*(2566) = - 20.15, *p* = 5.96×10^−84^, Cohen’s *d* = -0.834; **Fig. 1E**). Although additional differences between behavior of amygdala- and VS-lesioned monkeys have been reported previously^14^, better performance of VS-compared to amygdala-lesioned monkeys (two-sample t-test: *t*(2431) = - 12.35, *p* = 5.07×10^−34^, Cohen’s *d* = -0.519; **Fig. 1E** inset) is surprising considering the established role of VS for stimulus-based learning^24,25^ required for the What-only task.

During What blocks of What/Where task, however, amygdala-lesioned monkeys performed significantly better than VS-lesioned monkeys (two-sample t-test: *t*(1832) = 17.72, *p* = 6.05×10^−65^, Cohen’s *d* = 0.83; **Fig. 1F** inset), while both lesioned groups were impaired relative to controls (amygdala: *t*(3990) = 19.10, *p* = 1.01×10^−77^, Cohen’s *d* = 0.7; VS: *t*(3854) = 34.6, *p* = 1.08×10^−228^, Cohen’s *d* = 1.34). In Where blocks that did not require stimulus-based learning, only amygdala-lesioned monkeys showed impairments in performance relative to controls (two-sample t-test: *t*(3816) = -8.91, *p* = 8.07×10^−19^, Cohen’s *d* = -0.342; **Fig. 1G** inset). VS-lesioned performance was comparable to that of controls (two-sample t-test: *t*(3780) = -2.17, *p* = .0304, Cohen’s *d* = -0.08454) and better than that of amygdala-lesioned monkeys (*t*(3780) = 5.95, *p* = 3.31×10^−9^, Cohen’s *d* = 0.286), suggesting no deficits for action-based learning in the VS lesion group.

Overall, these results demonstrate that in the absence of uncertainty about the correct model of the environment and when this model is stimulus based, VS-lesioned monkeys (with intact amygdala), and not amygdala-lesioned monkeys, were able to partially overcome the deficit in stimulus-based learning. Under uncertainty about the correct model of the environment (What/Where), however, VS lesions caused significant impairment in What blocks only, consistent with the role of VS in stimulus-based learning. In contrast, amygdala-lesioned monkeys exhibited impaired performance in both What and Where blocks (but more strongly in What blocks) despite no clear evidence for the significant contribution of the amygdala to action-based learning (but see^17^), whereas there are action encoding in the amygdala and VS^26,27^. The similar impairments observed in amygdala-lesioned monkeys during What and Where blocks cannot be explained by the amygdala’s currently assumed role in stimulus-based learning.

These results are also puzzling because higher performance of VS-lesioned compared to amygdala-lesioned monkeys in the What-only task could be seen as evidence for stronger contribution of the amygdala to stimulus-based learning, yet higher performance of amygdala-lesioned compared to VS-lesioned monkeys in the What/Where task contradicts this notion. These findings hint at the potential role of the amygdala in arbitration between stimulus- and action-based learning in addition to its role in stimulus-based learning.

To study the relative adoption of the two learning strategies according to uncertainty of the reward environment, we examined the difference between ERDS_Stim_ and ERDS_Action_ by block types and reward schedules (**Supplementary Fig. S1**). We found that larger reward uncertainty leads to increasing influence of the competing (incorrect) strategy for respective block types across all groups. More specifically, in What-only and What blocks of the What/Where task, animals’ strategies became more action-based as the uncertainty in reward schedules increased (**Supplementary Fig. S1D, E**). Consistently, in Where blocks, they tended to become more stimulus-based under more uncertainty (**Supplementary Fig. S1F**). These observations demonstrate that both control and lesioned monkeys adjust similarly to reward uncertainty by shifting their strategies away from the more uncertain model, even though they start from different baselines. Critically, amygdala-lesioned monkeys exhibited a least overall distinction between the two types of learning strategies (**Supplementary Fig. S1D–F**).

Finally, we also examined reaction time in the two lesioned groups and found consistent results to those of the control animals (**Supplementary Note 1**). Together, our findings suggest that VS lesions biased behavior toward action-based learning by impairing stimulus-based learning. In contrast, amygdala lesions caused more nuanced impairment to both stimulus- and action-based learning and their coordination thereof. To reveal the underlying mechanisms, we next examined fit of choice behavior in control and lesioned monkeys.

### Mechanisms of arbitration between stimulus- and action-based learning systems

To uncover mechanisms underlying the interaction between the two systems, we developed multiple reinforcement learning (RL) models to fit choice behavior of control monkeys on a trial-by-trial basis. In the simplest model, signals from distinct action-based and stimulus-based learning systems were added linearly (weighted by a fixed parameter, *ω*) to determine the final decision variable. We also tested models with dynamic arbitration in which *ω* was adjusted on a trial-by-trial basis using the reliability of the two systems. Based on previous literature, we compared two ways by which the reliability could be computed: (1) magnitude of the reward prediction error (|RPE|); and (2) value of the chosen option (V_cho_). Additionally, we considered a more general model in which the baseline ratio of value signals from the two learning systems (quantified by parameter, *ρ*) could be adjusted independently of *ω* (**Fig. 2A;** see **Methods** for more details). As a result, this (Dynamic *ω*-*ρ*) model relaxes the assumption that the amount of increase in the strength of signals from one system (or equivalently, the sensitivity of final choice to those signals) is identical to the decrease in the strength of signals from the other system, and vice versa. To determine the best model, we computed the goodness-of-fit using five-fold cross-validation.

**Fig. 2.**
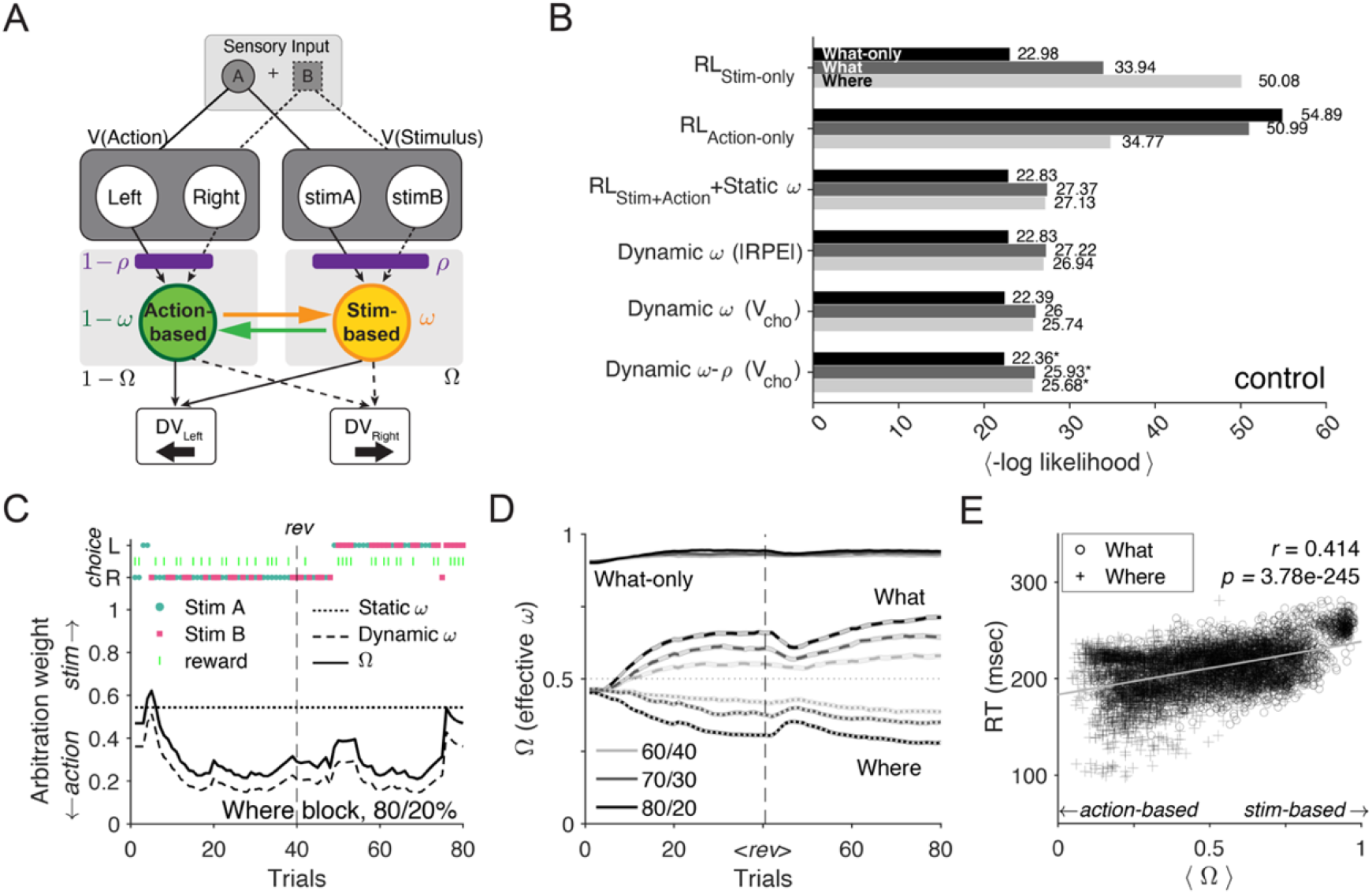
Fit of choice behavior of control monkeys using multiple RL models. (**A**) Schematic of the RL model with two parallel learning systems, showing an example trial in which the stimulus A appeared on the left side. In the static model, a constant *ω* is assumed to be fixed for each block of trials. In the dynamic models, *ω* is updated on each trial according to relative reliability of the two systems. In a more general dynamic model, the fixed parameter *ρ* (estimated for each subject) adjusts the baseline ratio of two value signals. For *ρ* = 0.5, the Dynamic *ω*-*ρ* model reduces to the Dynamic *ω* model. (**B**) Comparison of goodness-of-fit across models. Plotted is the mean negative log-likelihood over all cross-validation instances, separately for each task and corresponding block types: What-only (black), What (dark gray), Where (light gray). (**C**) Example Where block in the What/Where task and estimated arbitration weight from the Static *ω* model (dotted line), and arbitration weights (*ω*, dashed line) and effective arbitration weights (*Ω*, solid line) from the best model (Dynamic *ω*-*ρ* model). In this example, *ρ* = 0.61, effectively biasing behavior toward stimulus-based strategy. In this block, rightward action (R) was a better option than leftward action (L) before reversal (*rev*, horizontal dashed line). (**D**) Average trajectory of *Ω* from the Dynamic *ω*-*ρ* model during different tasks and blocks: What-only (solid), What (dashed), and Where (dotted). Different colors correspond to different reward schedules: 80/20 (black), 70/30 (dark gray), 60/40 (light gray). <*rev*> indicates reversal (horizontal dashed line), of which positions are normalized across blocks. (**E**) Relationship between *Ω* (block-averaged) and median reaction time for a given block during the What/Where task. Reported are Spearman’s correlation coefficient *r* and its p-value for all blocks during the What/Where task.

Comparing the single-system models with the simplest two-system model that assumes a fixed relative weighting for the two systems (RL_Stim+Action_ +Static *ω*), we found the latter to provide a better fit. Interestingly, this model improved the goodness-of-fit even in the What-only task in which action-learning was not predictive of reward. Overall, however, all the dynamic models provided better fit than the model with fixed weighting. Ultimately, the dynamic model that used the value of the chosen option (V_cho_) to estimate reliability in addition to a baseline weighting of two systems quantified by *ρ*, the Dynamic *ω*-*ρ* model, provided the best fit across all tasks (**Fig. 2B**).

To gain more insight into how dynamic arbitration improves the fit of choice behavior, we next examined the behavior of the Dynamic *ω*-*ρ* model and its arbitration weights over time. To that end, we computed the effective arbitration weight (“effective” *ω* denoted by *Ω*) to measure the overall relative weighting between two systems considering *ρ* (see **Methods** for more details). Both the example block and the averaged trajectories of trial-by-trial *Ω* from the best model (**Fig. 2C, D**) showed dependence on the block type and uncertainty in the reward schedule, especially during the What/Where task that required arbitration between competing models of the environment. These results demonstrate that *Ω* can capture behavioral adjustments to uncertainty over time.

We also tested whether this model could capture the difference in RT according to the more dominant system identified with ERDS, where we found that stimulus-based choices lead to slower RTs (**Supplementary Note 1**). To that end, we computed the correlation between the median RTs of the block and the average estimated values of *Ω*, which measures the overall relative weighting of the stimulus-based to action-based system. For What-only task, we found a small yet significant correlation between the effective weighting and RT (Spearman’s *r* = .094, *p* = 1.24×10^−4^). In comparison, *Ω* and RT were highly correlated in the What-Where task (*r* = .414, *p* = 3.78×10^−245^; **Fig. 2E**). This means that slower (respectively, faster) RT occurred when larger weights were assigned to the stimulus-based (respectively, action-based) system, consistent with the previous analysis in which action-dominant trials based on ERDS accompanied by faster RT.

As part of our exploration of arbitration mechanisms, we also compared multiple algorithms for estimating reliability of the two systems, including V_cho_, |RPE|, discernibility between two competing options (|ΔV|), and the sum of value estimates within each system (ΣV) (see **Methods** for details). We found that among the four reliability measures considered, V_cho_ best explained the control monkeys’ choice behavior across all block types (**Supplementary Fig. S2A)**. Specifically, the Dynamic *ω* model based on V_cho_ improved the fit over the Static *ω* model in all tasks, whereas the Dynamic *ω* model based on |RPE| improved the fit over the Static *ω* model only in the What/Where task (**Fig. 2B**). Through model recovery, we confirmed that our model fitting procedure could effectively discriminate between the alternative one-system and two-system models (**Supplementary Fig. S3A–D**).

Importantly, V_cho_ and |RPE| are conceptually similar as both signals measure the predictiveness of reward value in each system. However, the two signals differ in their sensitivity to negative feedback. For example, when the chosen value of the more reliable system (e.g., stimulus-based system in What block) is (correctly) estimated to be high, negative RPE (and consequently |RPE|) will also be high, undesirably facilitating the update toward the incorrect system. In comparison, V_cho_ by itself is less sensitive to negative feedback, as the updated value of V_cho_ after omission of reward will still reflect the high value estimates for the more reliable system. As a result, V_cho_ signal distinguishes the more reliable system better than |RPE| signal especially for more uncertain reward schedules (**Supplementary Fig. S2B–D**). Ultimately, difference in the V_cho_ drives the arbitration process (**Eqs. 8-9**), and this difference is equal to the difference in *signed* RPE. This suggests that the reliability signal could be more connected to RPE than unsigned RPE.

Finally, to further validate our models, we compared the predictions of different models regarding the observed negative interaction between ERDS_Stim_ and ERDS_Action_ during the What-only task. This is to ensure that this relationship was due to competition between the two learning systems and not due to task structure, as the animals could not stay/switch on the stimuli and location dimension at the same time while positions of stimuli were pseudo-randomly assigned to either side. To that end, we simulated choice behavior using single-system or two-system models, and computed regression weights between ERDS_Stim_ and ERDS_Action_ (see **Methods** for more details). Competition between the two learning systems during the What-only task would suggest that weaker stimulus-based learning corresponds to stronger action-based learning and vice versa. We found that in the single-system model that learned the stimulus-outcome contingencies only (RL_Stim-Only_), ERDS_Stim_ was only weakly predictive of ERDS_Action_ (**Supplementary Fig. S4B**; *β* = -0.006, *p* = 8.1×10^−45^). In contrast, in both the static and dynamic two-system models, ERDS_Stim_ was negatively predictive of ERDS_Action_, thus reproducing the competitive interaction between the two systems (**Supplementary Fig. S4C**; *β* = -0.038, *p* = 3.7×10^−28^; **Supplementary Fig. S4D**; *β* = -0.054, *p* = 7.3×10^−29^). These simulation results further corroborate the presence of multiple learning systems, even in an environment where one of the systems was not beneficial for performing the task.

### Deficits in arbitration due to amygdala but not VS lesions

Fit of choice behavior of lesioned monkeys revealed that the Static *ω* model explained the choice behavior of both lesioned groups better than models with a single learning system (**Fig. 3A**). Moreover, adding dynamic arbitration as in the Dynamic *ω* model further improved the fit from the Static *ω* model in both groups, and including the baseline relative weighting (Dynamic *ω*-*ρ* model) resulted in the best overall fit (**Fig. 3A**). Finally, consistent with the results in controls, for both lesioned groups, the dynamic model with V_cho_ as the basis for reliability signal accounted for choice behavior better than the model with |RPE| for reliability (**Supplementary Fig. S2A**).

**Fig. 3.**
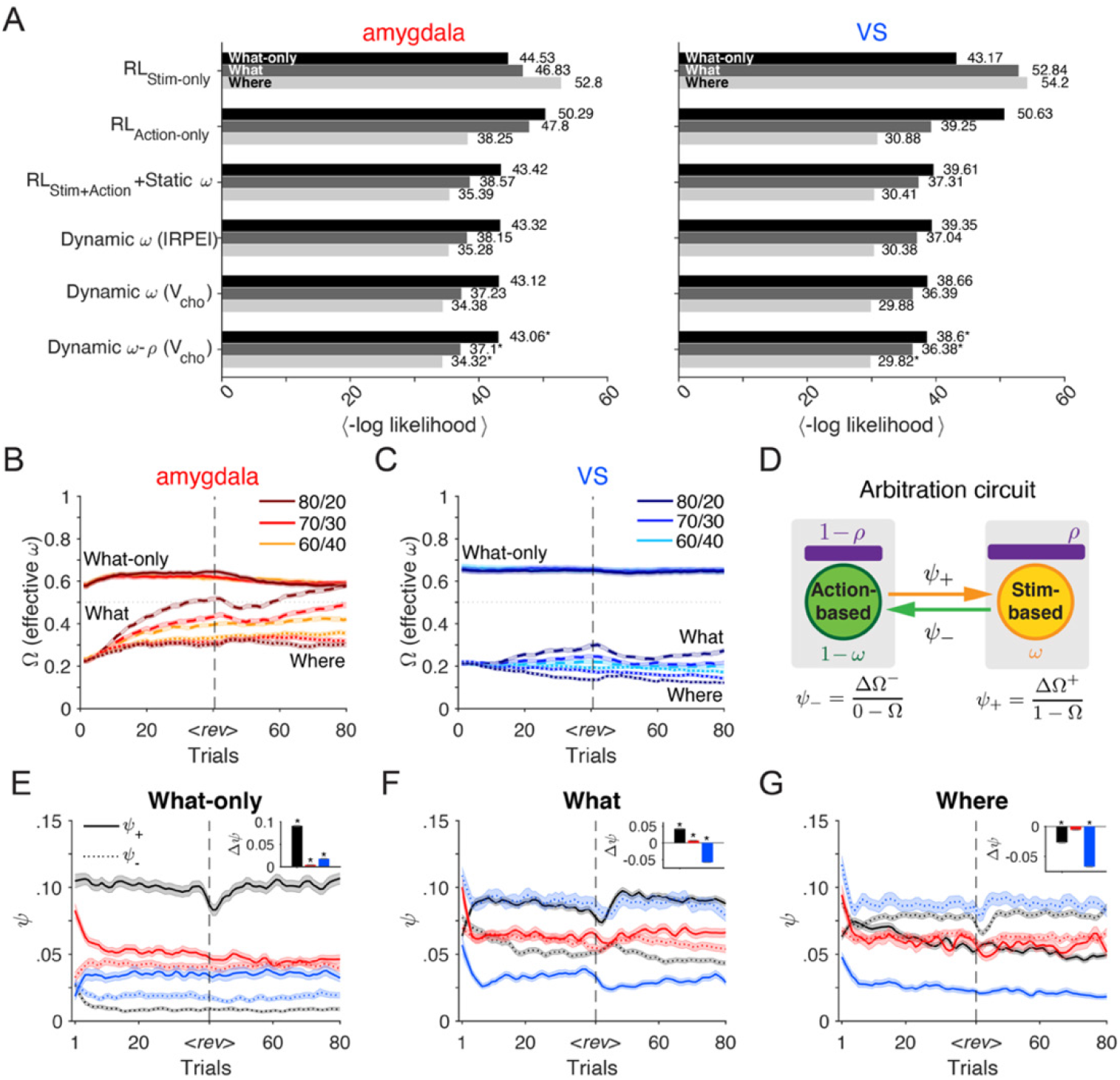
Fit of choice behavior for amygdala- and VS-lesioned monkeys. (**A**) Comparison of the models’ goodness-of-fit for choice behavior of amygdala-(left) and VS-lesioned (right) monkeys. Plotted is the mean negative log likelihood over all cross-validation instances for respective block types: What-only (black), What (dark gray), and Where (light gray). (**B–C**) Averaged trajectory of estimated *Ω* (effective *ω*) in amygdala-lesioned (B) and VS-lesioned (C) monkeys, separately for each block type and reward schedule. Solid, dashed, and dotted curves indicate What-only, What, and Where blocks, respectively. <*rev*> indicates reversal (horizontal dashed line), normalized across blocks. Colors indicate different reward schedules: 80/20 (brown and navy), 70/30 (red and blue), 60/40 (orange and cyan). (**D**) Schematic of effective arbitration rates in the arbitration circuit. *ψ*^+^ and *ψ*^−^ represent the rate of update toward the stimulus-based (increase in *Ω*) and action-based system (decrease in *Ω*). (**E–G**) Plotted is the time course of effective arbitration rates toward the stimulus-based or action-based system separately for each task (indicated on the top) and for each group of monkeys (controls: black; VS-lesioned: blue; amygdala-lesioned: red). Solid and dotted lines indicate effective arbitration rates toward stimulus-based (*ψ*^+^) and action-based systems (*ψ*^−^), respectively. Insets show the mean paired differences between the two arbitration rates within each block after reversal (Δ*ψ* = *ψ*^+^ − *ψ*^−^).

To determine the mechanisms by which different lesions impact the arbitration process, we examined the estimated parameters in the best model. We first confirmed that the parameters of this model were recovered well (**Supplementary Fig. S3E, F**). Estimated trajectory of *Ω* revealed that similar to controls, arbitration was modulated by reward uncertainty during the What/Where task in both amygdala-(**Fig. 3B**) and VS-lesioned monkeys (**Fig. 3C**). Nonetheless, *Ω* values were overall smaller than in controls, corresponding to more action-based strategy in lesioned animals (compare **Fig. 3B, C** and **Fig. 2D**). Importantly, a key difference between the two lesioned groups demonstrates deficits in the arbitration process due to amygdala lesions. Specifically, in the What-only task (solid curves in **Fig. 3B**), *Ω* was stable during initial learning but slightly decreased toward 0.5 after reversals, whereas it increased over time during both What and Where blocks (dashed and dotted curves in **Fig. 3B**). In contrast, VS-lesioned monkeys showed an increase in *Ω* for What blocks and a decrease in *Ω* for Where blocks (dashed and dotted curves in **Fig. 3C**) similar to control animals, while having a stable *Ω* during the What-only task (solid curves in **Fig. 3C**).

These results demonstrate that VS lesions biased behavior toward action-based learning while keeping the arbitration processes intact, whereas amygdala lesions impaired arbitration in addition to biasing behavior toward action-based learning. These suggest that the deficits observed in amygdala-lesioned monkeys cannot solely be attributed to impairments in stimulus-based learning; instead, they involve a more complex interaction and dynamic compensations between stimulus- and action-based signals. Consistent with this interpretation, we found that the winning model with dynamic arbitration (Dynamic *ω*-*ρ*) better captures the key aspects of behavioral strategy in the two lesioned groups when compared to the single-system model (**Supplementary Fig. S5**).

To further investigate the dynamics of the arbitration weight, we next examined the rate of changes in *Ω* across the three groups. To that end, we calculated the “effective” arbitration rates by computing the ratio of the overall change in *Ω* toward 1 (stimulus-based system) or 0 (action-based system) relative to its original value (**Fig. 3D**; see **Methods** for more details). This quantity measures the rate of arbitration, analogous to the learning rate for updating value estimates. We found that in control monkeys, the effective arbitration rates toward the stimulus-based (*ψ*_+_) or action-based (*ψ*_−_) system diverged soon after the beginning of a block, reflecting the correct model of the environment. When the stimulus-based system was more reliable, the effective arbitration rates toward the stimulus-based system were larger (What-only task, paired-samples t-test: *t*(1657) = 42.8, *p* = 3.50×10^−270^, Cohen’s *d* = 1.05; What/Where task, *t*(2986) = 36.0, *p* = 2.38×10^−236^, Cohen’s *d* = 0.663; **Fig. 3E, F**). Similarly, in Where blocks in which the action-based system was more reliable, the effective arbitration rate toward the action-based system was significantly larger than that toward the stimulus-based system (*t*(2928) = 21.6, *p* = 4.38×10^−96^, Cohen’s *d* = 0.402; **Fig. 3G**).

In contrast, amygdala-lesioned monkeys showed the minimum differentiation between adjustments toward the more and less reliable (correct and incorrect) systems. Specifically, during the What-only task, the difference between the two arbitration rates were significantly smaller for amygdala-lesioned compared to VS-lesioned monkeys (two-sample t-test: *t*(2401) = -2.71, *p* = .0068; Cohen’s *d* = -0.114; **Fig. 3E**). During the What/Where task, amygdala-lesioned monkeys exhibited a slight preference for transition toward the stimulus-based system in What blocks yet less strongly than controls (*t*(984) = 10.27, *p* = 1.41×10^−23^, Cohen’s *d* = 0.327; **Fig 3F**), while showing no significant difference between two arbitration rates during Where blocks (*t*(880) = -1.24, *p* = .214, Cohen’s *d* = -0.042; **Fig. 3G**). VS-lesioned monkeys, however, exhibited overall large bias in arbitration rates toward the action-based system during What/Where task in both blocks (What: *t*(848) = 30.6, *p* = 3.76×10^−139^, Cohen’s *d* = 1.05; Where: *t*(844) = 34.5, *p* = 2.30×10^−163^, Cohen’s *d* = 1.19; **Fig. 3F**,**G**). In the What-only task, VS-lesioned group maintained higher arbitration rates toward the stimulus-based system, thus appropriately biasing the correct system for the task (*t*(893) = 9.55, *p* = 1.17×10^−20^; Cohen’s *d* = 0.32; **Fig. 3E**). Together, these results suggest that amygdala lesions impair arbitration between the two learning systems by eliminating differential updates for the correct and incorrect (more and less reliable) systems, thus hinting that amygdala is crucial for identifying the correct model of the environment. In contrast, VS lesions mainly impair stimulus-based learning and overall arbitration bias toward action-based learning but only under uncertainty about the correct model of the environment (What/Where).

### Dynamic interaction between learning and arbitration processes and the impact of initial state

Considering the observed effects of amygdala and VS lesions on arbitration dynamics, we next examined the estimated parameters from the best-fit model (Dynamic *ω*-*ρ*). In this model, *ρ* captures whether there is an overall reduction in signal from stimulus-based system relative to action-based one. We found that, for both What-only and What/Where tasks, the estimated values of *ρ* were smaller in VS-than amygdala-lesioned monkeys, indicating a larger baseline reduction in stimulus-based signals relative to action-based signals in VS-lesioned monkeys (**Fig. 4A**). As a result, VS-lesioned monkeys exhibited larger sensitivity of choice to action-based signals (*β*_action_) than to stimulus-based signals (*β*_stim_) during both tasks (paired-sample t-test; What-only: *t*(83) = 3.43, *p* = 8.89×10^−4^, Cohen’s *d* = 0.376; What/Where tasks: *t*(89) = 16.7, *p* = 3.14×10^−29^, Cohen’s *d* = 1.76; **Fig. 4B, C**, blue data points). This was not the case for the amygdala-lesioned monkeys which showed significantly larger *β*_stim_ than *β*_action_ in both tasks (What-only: *t*(117) = 8.1, *p* = 5.91×10^−13^, Cohen’s *d* = 0.746; What/Where: *t*(130) = 12.1, *p* = 5.56×10^−23^, Cohen’s *d* = 1.06; **Fig. 4B, C**, red data points). This was also the case for control monkeys (What-only: *t*(91) = 14.1, *p* = 1.20×10^−24^, Cohen’s *d* = 1.47; What/Where: *t*(310) = 15.4, *p* = 5.59×10^−40^, Cohen’s *d* = 0.871; **Fig. 4B, C**, black data points). Interestingly, the differences between sensitivity of choice to two signals (Δ*β*) were not significantly different between control and amygdala-lesioned monkeys in both What-only (two-sample t-test: *t*(208) = 0.906, *p* = .366, Cohen’s *d* = 0.126; **Fig. 4B** inset) and What/Where tasks (*t*(440) = 0.396, *p* = .692, Cohen’s *d* = 0.041; **Fig. 4C** inset). These results suggest that unlike VS lesions, amygdala lesions did not affect the relative preference for stimulus-based vs. action-based signals. Therefore, in line with previous observations, the deficit observed from amygdala lesions cannot be solely attributed to impairments in the stimulus-based learning. Instead, they suggest deficits in arbitration processes that subsequently affect learning processes.

**Fig. 4.**
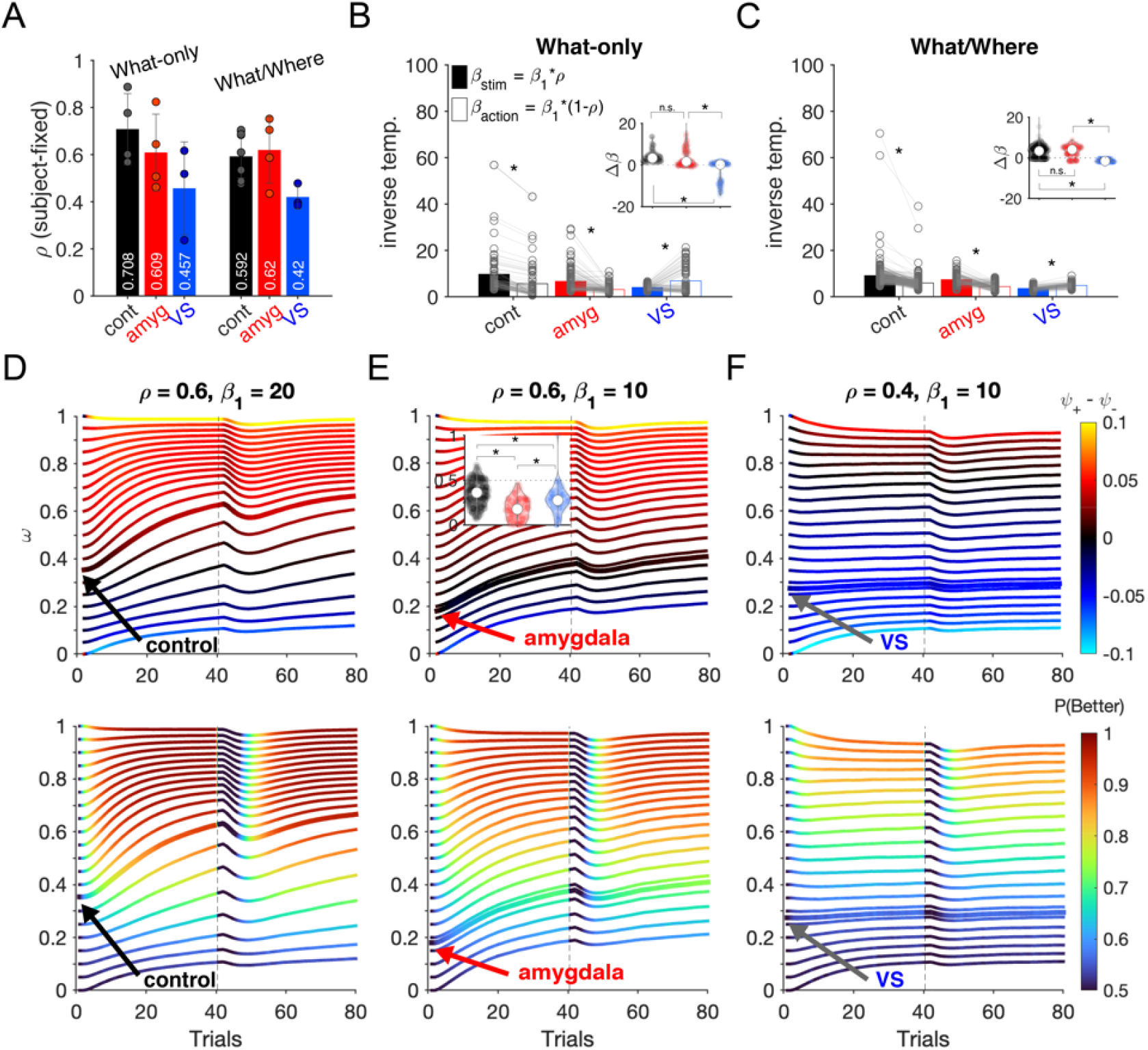
Comparison of relative weighting and sensitivity of choice to stimulus- and action-based signals across control and lesion groups, and their simulations. (**A**) Plots show the mean and individual values (each point represents a monkey) of the relative strength of two systems on choice (*ρ*), separately for each group and task. (**B–C**) Comparison of sensitivity of choice to signals in the two systems, separately for each group and task. *β*_1_ is the common sensitivity of choice (inverse temperature) estimated for each session and *ρ* is the relative strength of the two systems as in panel A. Asterisks indicate significant difference between sensitivity to two systems by paired-sample t-test (*p* < .001). Insets are the violin plots of paired differences (Δ*β* = *β*_stim_ – *β*_action_), and asterisks indicate significant between-group differences by two-sample t-test (*p* < .001). Only VS-lesioned monkeys showed larger sensitivity to action-than stimulus-based systems during both tasks, consistent with the role of the VS in stimulus-based learning. (**D–F**) Plots show simulated trajectory of *ω* and difference in the effective arbitration rates (*ψ*_+_ – *ψ*_-_; top) and performance (P(Better), bottom). Each line represents averaged trajectories (10,000 simulated blocks) of *ω* with different initial values (*ω*_0_) and specified values of *ρ* and *β*_1_ during the What task. All other parameters are fixed (*α*_+_ = *α*_-_ = 0.5, *β*_0_ = 0, ζ= 0.3, *α*_*ω*_ = 0.2, ζ_*ω*_ = 0.05). Black, red, and gray arrows show the trajectory simulated with *ω*_0_ equal to the mean of control, amygdala-lesioned, and VS-lesioned monkeys, showing *ψ*_+_ > *ψ*_-_, *ψ*_+_ ≈ *ψ*_-_, and *ψ*_+_ < *ψ*_-_, respectively. Horizontal line (trial 40) indicates reversal. Insets show distribution of *ω*_0_ across the three groups during the What/Where task and asterisks indicate significant difference by Wilcoxon rank sum test (*p* < .001).

To confirm this point, we examined the initial arbitration weights that determine the weights of the two systems on choice at the beginning of each block, when the monkeys are oblivious to the correct model of the environment during What/Where task. We note that amygdala- and VS-lesioned monkeys did not significantly differ in the initial *Ω* (effective *ω*) values (rank sum test: *p* = 0.0926; compare *Ω* of the first trial in **Fig. 3B** and **Fig. 3C**). However, by examining the baseline arbitration weights before scaling by *ρ*, we found that *ω*_0_ values were significantly smaller in amygdala-compared to VS-lesioned monkeys (rank sum test: *p* = 5.05e×10^−7^; **Fig. 4E** inset). This means that larger values of *ρ* in amygdala-lesioned monkeys were offset by lower *ω*_0_ values to yield *Ω*_0_ comparable to VS-lesioned monkeys. In other words, deficits due to amygdala lesions can be attributed mainly to *ω*_0_ and subsequent interaction between arbitration and learning processes, whereas deficits due to VS lesions are largely caused by baseline reduction in stimulus-based signals, measured by *ρ*.

To further support this idea, we simulated the choice behavior of the Dynamic *ω*-*ρ* model by adjusting two key parameters: baseline ratio of learning signals, *ρ*, and the initial arbitration weight, *ω*_0_. We kept all other parameters constant except for the common inverse temperature, *β*_1_. Trajectories of simulated *ω* during What blocks revealed that different values of *ρ* and *ω*_0_ can create different dynamics with respect to arbitration rates (**Fig. 4D–F**, top). More specifically, the simulated trajectory of *ω* based on mean *ω*_0_ in control monkeys during the What/Where task (**Fig. 4D**, black arrow) resulted in larger transitions toward the stimulus-based system (*ψ*_+_ > *ψ*_-_), whereas mean *ω*_0_ in amygdala-lesioned monkeys (**Fig. 4E**, red arrow) reduced the distinction between the two arbitration rates (*ψ*_+_ ≈ *ψ*_-_). In comparison, simulations using mean *ω*_0_ in VS-lesioned monkeys (**Fig. 4F**, gray arrow) resulted in preferred update toward the action-based system (*ψ*_+_ < *ψ*_-_). These results mimic the pattern of arbitration rates in the three groups.

Finally, we also tested the causal contribution of initial arbitration weight on performance using simulated choice behavior (**Fig. 4D–F**, bottom). Crucially, we found that lower values of *ω*_0_, as observed in amygdala-lesioned monkeys (*ω*_0_ = 0.18; red arrows in **Fig. 4E**, bottom), lead to reduced performance when compared to higher *ω*_0_ values (e.g., *ω*_0_ = 0.40). This effect is reflected in a significant main effect of *ω*_0_ on the simulated performance (*F*(20,1659) = 40.6, *p* = 8.38×10^−128^). These simulation results demonstrate that inflexibility in the arbitration process, as modeled by a reduction in *ω*_0_, could be the primary cause of impaired performance, rather than a secondary effect.

Overall, these findings also suggest that the initial state of the system (*ω*_0_) is crucial for determining the later trajectory and rates of transition in the arbitration process. Our simulations show that amygdala-lesioned monkeys occupy a parameter space that yields less distinction between arbitration rates toward and away from the correct system in the given environment, ultimately resulting in reduced performance. Although control monkeys were also biased toward the action-based system at the start of the What/Where task (mean *ω*_0_ ∼ 0.36), lesions to the amygdala resulted in an even larger bias toward the action-based system (mean *ω*_0_ ∼ 0.18), and this consequently led to lack of differential updates for the two systems. In contrast, lesions to VS mainly decreased *ρ* to bias signals toward the action-based system, while affecting the initial state of arbitration to a lesser degree (mean *ω*_0_ ∼ 0.27).

### Diversity of behavior driven by dynamic interaction between learning and arbitration processes

To demonstrate the impact of dynamic interaction between the learning and arbitration processes, we simulated the model within the task by adjusting parameters such as the learning and forgetting rates. These simulations revealed a wide range of dynamics in performance and arbitration weights, highlighting complex interactions between learning, arbitration, and decision-making processes (**Fig. 5**). Interestingly, we observed that higher initial arbitration weights, which would allow the animals to correctly bias their behavior toward the stimulus-based system during a stimulus-learning task, can both facilitate and impede learning after reversals depending on other parameters of the model. That is, in most cases, larger initial bias toward the stimulus-based system helps both initial learning of stimuli and their reversals (**Fig. 5A**,**C**,**D**,**F**).

**Fig. 5.**
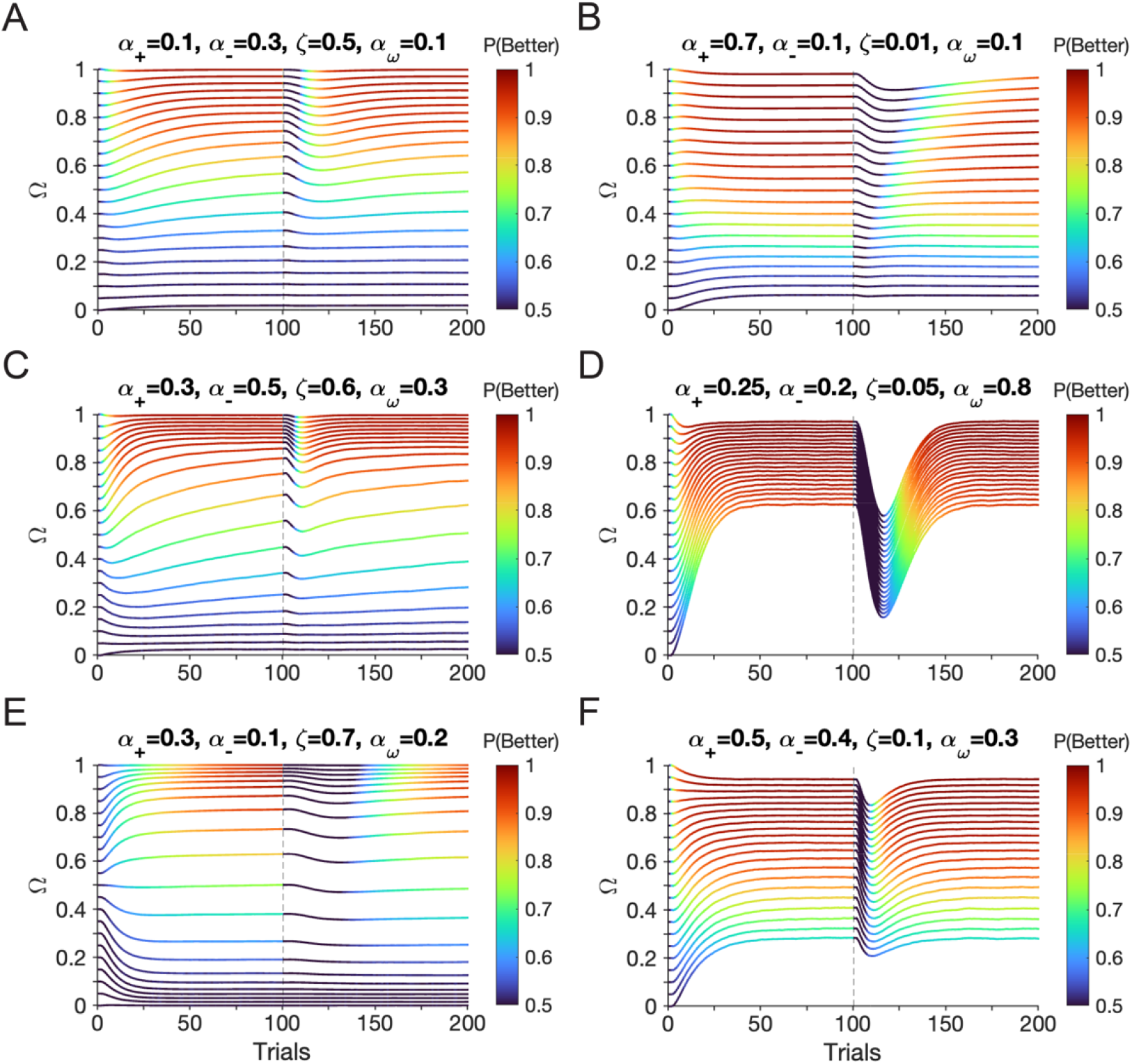
Complex interaction between arbitration and learning creates diverse learning patterns. Each line represents averaged trajectories (10,000 simulated blocks) of *Ω* with different initial values during a stimulus-based learning task with reversal at trial 100. All simulations were performed with *ρ* = 0.5, causing *Ω* = *ω*. Panels A–F show different behavioral patterns based on simulation of choice behavior using different model parameters, as indicated on the top. All non-specified parameters are fixed across panels at *β*_1_ = 20, *β*_0_ = 0, and ζ_*ω*_ = 0.05. (**A**) Larger initial values (*ω*_0_) facilitate learning after reversal. (**B**) Larger initial values (*ω*_0_) impede learning after reversal. (**C**) Initial values *ω*_0_ > 0.15 increase stable points of *ω* toward 1, whereas small *ω*_0_ (< 0.15) results in low performance. (**D**) Small decay or forgetting for unchosen option (ζ) and large transition rate (*α*_*ω*_) facilitates arbitration toward the correct model. (**E**) Bifurcation of trajectories happens around *ω* of 0.5. (**F**) Steady state of arbitration is controlled by the initial value.

However, in scenarios where the positive learning rate significantly exceeds the negative learning rate, a smaller initial arbitration weight—though it may incorrectly bias behavior toward an action-based strategy—can actually facilitate adjustments to reversals in stimulus values (**Fig. 5B**). This happens because lower values of *ω*_0_, as in the case of amygdala lesions, result in a more mixed dependence of choice on stimulus- and action-based signals and thus, more explorations that greatly benefit response to reversals. These results, based on our dynamic arbitration model, can thus explain the paradoxical improvements in performance observed following amygdala lesions or inactivation.

### Contribution of amygdala to long-term adjustments of behavior

Lesions to certain brain areas are often accompanied by adjustments or compensation by other brain areas that result in reducing initial behavioral impairments over the long term. Considering the observed effects of amygdala and VS lesions on learning and decision-making behavior, we investigated long-term adjustments in these behaviors in the absence of uncertainty about the correct model of environment. To that end, we examined ERDS and median RT across all sessions of the What-only task using the proportion of sessions completed as an independent variable.

For consistent use of stimulus-based strategy, we observed a significant decrease in ERDS_Stim_ in VS-lesioned monkeys (VS: *β* = -0.246, *p* = 1.52×10^−15^), whereas there was no evidence for changes in control or amygdala-lesioned monkeys across time (Control: *β* = -0.0479, *p* = .138; amygdala: *β* = -0.0416, *p* = .0748; **Supplementary Fig. S6A**). Specifically, despite their impaired stimulus-based learning, monkeys with VS lesions were able to increase their adoption of stimulus-based strategy over time. Consistently, VS-lesioned monkeys also decreased their adoption of action-based strategy as reflected in the positive slope of ERDS_Action_ over time (*β* = 0.106, *p* = .00133; **Supplementary Fig. S6B**). There was no evidence for such effect in control monkeys (*β* = -0.0139, *p* = .135) or in monkeys with amygdala lesions (*β* = 0.0299, *p* = .0676). Interestingly, consistent with previous results, the complementary changes in model adoption in VS-lesioned monkeys were also reflected in increased median RT over time in these monkeys (*β* = 14.997, *p* = 7.64×10^−7^; **Supplementary Fig. S6C**) but not in control monkeys (*β* = 3.98, *p* = .655) or amygdala-lesioned monkeys (*β* = 0.107, *p* = .985). This was accompanied by a long-term increase in the initial effective arbitration weights *Ω*_0_ (toward stimulus-based system) in VS-lesioned monkeys (*β* = 0.147, *p* = 8.76×10^−4^; **Supplementary Fig. S6D**).

These results provide evidence for adjustments on a long timescale in VS-but not amygdala-lesioned monkeys. They suggest that in the absence of uncertainty about the model of the environment, intact amygdala in VS-lesioned monkeys (and not intact VS in amygdala-lesioned monkeys) enabled these animals to slowly improve their stimulus-based learning over time. This amygdala-driven mechanism enabled VS-lesioned monkeys to gradually suppress action-based strategy, resulting in an increase in overall RT and estimated *Ω*_0_ over time.

## Discussion

Here, we applied a combination of approaches to examine data from control monkeys and monkeys with amygdala and VS lesions to explore the interaction between stimulus- and action-based learning and to reveal computational and neural mechanisms underlying arbitration processes. Using multiple behavioral metrics, we found strong evidence for competitive interaction between the two learning systems. Moreover, by constructing multiple network models and fitting choice data with these models, we tested the plausibility of different mechanisms for estimating reliability signals that guide arbitration processes. Our model simulations also elucidated how the interactions between learning, arbitration, and choice processes give rise to a complex dynamical system for learning behavior with strong dependency on the initial state.

Previous studies have identified deficits in both stimulus-based and action-based learning due to amygdala lesions^14,15^, but considered these deficits independently of each other. These studies concluded that amygdala lesions reduce choice consistency (sensitivity to value signals) for stimulus-based learning^14^ and increase sensitivity to negative feedback (*α*_-_) for action-based learning^15^. In addition to replicating these results (**Supplementary Fig. S7B, D**), our study provides a unified account by conceptualizing the effects of amygdala lesions in terms of arbitration between competing models of the environment under uncertainty.

Specifically, we found that the difference between *β*_stim_ and *β*_action_ in amygdala-lesioned monkeys were not significantly different from that of controls (**Fig. 4B, C** inset), suggesting that impairments in stimulus-based learning and action-based learning were comparable in amygdala-lesioned monkeys. Instead, the main deficits due to amygdala lesions can be captured with biased initial states of arbitration toward action-based signals, which consequently change the interaction between learning and arbitration, leading to impaired performance. When coupled with reduced choice consistency, this effect diminishes the distinction between the effective arbitration rates for correct and incorrect (more reliable and less reliable) models (**Fig. 4E**). This could imply that amygdala is important for identifying the more reliable model of the environment^13^ or mediating the effect of such identification on arbitration processes. This suggests that the amygdala, like the prefrontal cortex, is involved in learning to learn ^28^ and can explain why amygdala lesions weaken the amount of evidence needed before the animals reverse their choice preference^13^.

In comparison, we observed that VS-lesioned monkeys (with intact amygdala) were able to gradually overcome their impaired stimulus-based learning while showing a significantly larger arbitration rate for the stimulus-based system during the What-only task (**Fig. 3E**). This suggests that a signal to or from the amygdala, but not in the amygdala-to-VS pathway, could bias arbitration toward the more reliable model and lead to slow long-term behavioral adjustments. We found that arbitration was still present in amygdala-lesioned monkeys, suggesting that the amygdala is not required for arbitration per se but has a more nuanced role by setting the initial balance between models and improving the overall sensitivity to value signals. These two effects result in larger arbitration rates for the more reliable model of the environment, thus altering the trajectory of learning and choice behavior.

Arbitration between alternative models has garnered significant interest in cognitive, behavioral, and systems neuroscience. This includes arbitration between model-free vs. model-based RL^29– 32^, Pavlovian vs. instrumental control^33^, habitual vs. goal-directed system^34,35^, competing sets of strategies for solving complex stimulus-response mappings^36^, and during social decision-making^37–39^. Here, we explored more basic arbitration required for any type of decision making, as any choice option has to be selected by taking an action. Unlike arbitration among different types of learning systems that requires different types of reliability signals (e.g., model-free vs. model-based, based on unsigned reward prediction error and unsigned state prediction error^30^), we found that the same reliability signal, based on the value of the chosen option or chosen action (V_cho_) can be used for arbitration. Critically, we found that in both controls and lesioned monkeys, the reliability signal based on V_cho_ captured arbitration better than the reliability signal based on |RPE|. Because reliability based on difference in chosen value is the same as reliability based on the difference in signed RPE, our results suggest that the reliability of alternative models could be more connected to RPE than unsigned RPE based on those models.

Our proposal for the amygdala’s contribution to stronger weighting of the signal from the more reliable model on arbitration process is consistent with its postulated role in signaling attentional shifts for relevant control of behavior^40,41^. There are several pathways through which the amygdala could affect model arbitration. One major candidate is prefrontal-amygdala circuits^42,43^. In particular, the orbitofrontal cortex (OFC) receives substantial projections from the amygdala^44,45^ and could serve a central role in encoding and monitoring the reliability of multiple actor predictive models^3^. Given that amygdala-to-OFC input has been reported to be significantly involved in value coding by OFC neurons^46,47^, it is possible that this input also carries information for selective arbitration to appropriately bias behavior toward the relevant learning system in a given environment. Conversely, the PFC-to-amygdala pathway could signal the internal state variable (arbitration weight in our model), serving as the necessary input to the amygdala for computing the differential adjustment that sends feedback to the PFC. This may explain why additional basolateral amygdala lesions could reduce OFC-induced impairment in reversal learning^11^. Thus, strong reciprocal connections between amygdala and PFC, in particular vlPFC, OFC, or ACC, could be crucial for proper arbitration between alternative models of the environment.

The role of the amygdala in instrumental learning has been a matter of debate due to mixed evidence both in favor of^14–16,48,49^ and against^11,12,50–52^ its involvement in RL. Critically, our framework can account for the amygdala’s seemingly inconsistent role. Through simulations of the model that best captures choice data, we found that in certain situations, the lower initial arbitration weight that biases behavior toward the action-based system can actually facilitate adjustments to reversals during stimulus-based learning. This happens because more reliance on the less reliable action-based system allows faster exploration of alternative stimuli and thus faster reversal. This can account for puzzling improvement in performance due to basolateral amygdala lesions in rats^12^ or monkeys^13,50^ which has been attributed to increased benefits from negative feedback. In these examples, the stimulus-based system would still prefer the previously better option, which is no longer rewarding, but a less reliable action-based system would cause more switching from that option. Other studies that reported null results from amygdala deactivations^51,52^ have also utilized object reversals after initial learning of stimulus-reward associations over a long period of time (referred to as discrimination learning).

What these studies have in common is that they all utilize visual discrimination learning over a long period of time before a reversal, which would suppress learning from unrewarded trials (i.e., small *α*_-_) and allow the reliability of the stimulus-based system to reach its asymptote, thus slowing down reversal. This is very different from our experimental paradigm in which reversals happened on a short timescale before the reliability of the stimulus-based system could stabilize. Overall, our study suggests that observed discrepancies in the effects of amygdala lesions on learning and choice behavior are due to a dynamic interaction between arbitration, learning, and choice processes.

Dynamic interaction between arbitration and learning processes is particularly relevant to experimental paradigms that assume the presence of one learning system while unintended competing systems could have strong influence. Our results indicate that interpreting behaviors shaped by various learning systems should be approached with caution. This is particularly important when the manipulations in use might influence the arbitration process and thereby change the interplay among the learning systems. In principle, a multitude of simple learning strategies could underlie the heterogeneity in the so-called decision variables^53–55^, and careful examination of neural signals^56–58^ is needed to properly identify the neural substrates of corresponding learning systems.

## Methods

### Experimental paradigm

We examined two variants of a probabilistic reversal learning task in which monkeys selected between two visual stimuli to obtain a juice reward. During each block of the What-only task, reward was assigned stochastically according to stimulus identity only while reward probabilities on the two stimuli (selected afresh on each block) switched between trial 30 and 50 of the block without any signal to the monkeys (**Fig. 1A**). In the What/Where task, reward was assigned based on either stimulus identity (What blocks) or stimulus location (Where blocks) with reversal similar to the What-only task (more details below). Data were collected from a total of seventeen monkeys, some of whom received bilateral excitotoxic lesions to either the amygdala or the ventral striatum (VS). All experimental procedures for all monkeys were performed in accordance with the *Guide for the Care and Use of Laboratory Animals* and were approved by the National Institute of Mental Health Animal Care and Use Committee. We describe each experimental setup in more detail below.

#### What-only task

Data from this task^14^ were collected in eleven male rhesus macaques weighing 6.5 to 10.5 kg (controls: *n* = 4; amygdala-lesioned: *n* = 4; VS-lesioned: *n* = 3). The monkeys completed an average of 26.73 sessions (SD = 5.98) and an average of 16.81 (SD = 6.72) blocks per session. Each block consisted of 80 trials and involved a single reversal of stimulus-outcome contingencies on a randomly selected trial between trials 30 and 50 from a uniform distribution (**Fig. 1A**). On each trial, monkeys were trained to fixate on a central point on a screen (500–750 ms) to initiate the trial. After fixation, two stimuli, a square and circle of random colors, were assigned pseudorandomly to the left and right of the fixation point (6° visual angle). Monkeys indicated their choice by making a saccade to the target stimulus and fixating for 500ms. Reward (0.085 ml juice) was delivered according to the assigned reward schedule for a given block. Each trial was followed by a fixed 1.5 s inter-trial interval. Trials in which monkeys failed to fixate within 5s or make a choice within 1s were aborted and then repeated.

The reward schedule was determined by the probabilities of reward on two choice options selected from four possible values: 100/0, 80/20, 70/30, and 60/40. The reward schedule was randomly selected at the start of each block and remained constant within that block. Monkeys performed the deterministic task (100/0 reward schedule) after the data collection for the stochastic task had been completed. Here, we focus on our analyses of the task’s stochastic variant to match the reward schedules for the What/Where task described below. All monkeys that received lesions were trained and tested following their recovery from surgery. For more detailed surgical information, see the Supplemental Experimental Procedures in^14^. This experimental setup and some analyses of the data have been previously reported^14^.

#### What/Where task

Unlike the What-only task, the What/Where task involved both stimulus-based and action-based learning, and this feature introduced additional uncertainty about the correct model of the environment. The effects of lesions to the VS and amygdala during the What/Where task were examined in two different studies with separate sets of controls for each. The first study investigating the effect of VS lesions^22^ had a total of eight subjects weighing 6.5– 11 kg (controls: *n* = 5; VS-lesioned: *n* = 3). Four of the five controls and all of the three VS-lesioned monkeys were the same monkeys used in the What-only task^14^. The second study investigating the effect of amygdala lesions^15^ had a total of ten subjects weighing 6–11 kg (controls: *n* = 6; amygdala-lesioned: *n* = 4). Four of the six unoperated controls were the same monkeys used in the What-only task ^14^ and the What/Where task involving VS lesion ^22^. One additional control monkey was used as an unoperated control for the earlier What/Where task only ^22^. The remaining control and any of the amygdala-lesioned monkeys were not used in any previous studies.

The monkeys in this task completed an average of 29.56 sessions (SD = 5.36), with an average of 18.41 (SD = 4.26) blocks per session. Each block consisted of 80 trials and involved a single reversal of the stimulus-based or action-based contingencies between trials 30 and 50. A given block was randomly assigned as What or Where block and remained constant within that block (**Fig. 1B**). In What blocks, reward probabilities were assigned based on stimulus identity, with a particular object having a higher reward probability. In Where blocks, reward probabilities were assigned based on location, with a particular side having a higher reward probability regardless of stimulus identity. What and Where blocks were randomly interleaved throughout the session and block type was not indicated to the monkey. The reward schedule was randomly selected from three schedules (80/20, 70/30, 60/40) at the start of each block and remained constant within that block.

On each trial, monkeys were trained to fixate on a central point on a screen (400–600ms) to initiate the trial. After fixation, two visual objects were assigned pseudorandomly to the left and right of the fixation point (6° visual angle). Each block used two novel images that the animal had never seen before. Monkeys indicated their choice by making a saccade to the target stimulus and fixating for 500ms. Reward was delivered probabilistically according to the assigned reward schedule for a given block. Each trial was followed by a fixed 1.5 s inter-trial interval. Trials in which monkeys failed to fixate within 5s or make a choice within 1s were aborted and then repeated. This experimental setup, surgical information, and some analyses of the data have been previously reported ^15,22^.

### Quantification and statistical analysis

#### Entropy-based metrics

Here, we utilized information-theoretic metrics to quantify learning and choice behavior^20,21^. This includes entropy of choice strategy (H(str)), mutual information of reward and strategy, and conditional entropy of reward-dependent strategy (ERDS). Out of these metrics, we focused on ERDS to demonstrate how monkeys linked reward feedback to stimulus identity and/or action.

Generally, ERDS is calculated as follows:

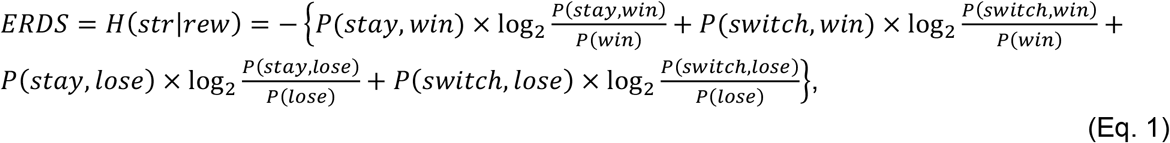

where *str* is the adopted strategy coded as stay (1) or switch (0), *rew* is the previous reward outcome coded as reward (1) or no reward (0). In this equation, 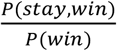 and 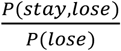 correspond to win-stay and lose-switch, respectively. Additionally, H(*str*) above is the entropy of strategy (stay or switch):

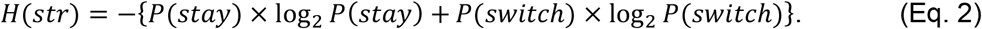

As the equations above suggest, ERDS measures the dependence of adopting a response strategy on reward feedback (*win* or *lose*). Lower ERDS values correspond to decreased randomness in the variable and thus more consistency in the utilized strategy, which could be stimulus-based and/or action-based. To detect these strategies, we defined two types of ERDS by considering choice and reward feedback in terms of stimulus identity or action, corresponding to ERDS_Stim_ and ERDS_Action_, respectively.

Therefore, lower values of ERDS_Stim_ suggest that the animals stay or switch consistently based on stimulus identity according to reward feedback, indicating the stronger adoption of the stimulus-based strategy. Conversely, lower values of ERDS_Action_ indicate that the animals adopted the action-based strategy more strongly. Overall, comparison of ERDS_Stim_ and ERDS_Action_ enables us to quantify the adopted strategy on a trial-by-trial basis, either by computing the average values across a block of trials or by aligning all trials relative to the beginning, reversal point, and the end of each block.

### Computational models

#### Single-system reinforcement learning (RL) models

We first used two standard RL models that learn about one type of reward contingencies to fit monkeys’ choice data. Specifically, the RL_Stim-Only_ and RL_Action-Only_ models associate reward outcomes to choice options either in terms of stimulus identity or chosen action in order to estimate stimulus and action values, respectively. These values were used to determine choice on each trial and were updated based on reward outcome at the end of trial, as described below.

More specifically, the value of the chosen option (*V*_*C*_) is updated using reward prediction error (RPE) and two separate learning rates for rewarded and unrewarded trials (*α*_+_ and *α*_-_, respectively) while the value of the unchosen option (*V*_*U*_) decays to zero:

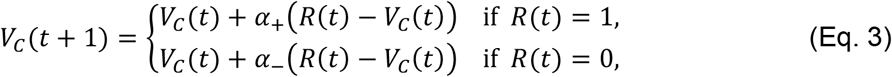

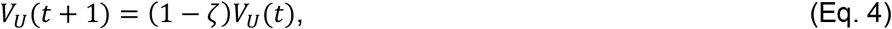

where *R*(t) is the reward feedback on trial *t* and ζ is the decay or forgetting rate for the unchosen option. Chosen and unchosen options are coded as {*stimulus A, stimulus B*} in the RL_Stim-Only_ model and as {*Left, Right*} in the RL_Action-Only_ model. In these and other models, the probability of choosing the option on the left, *P*_*left*_(*t*), was computed using a softmax function:

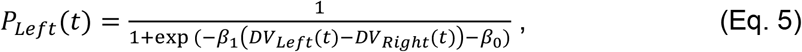

where *DV*_*left*_ and *DV*_*right*_ are decision variables for the options on the left and right, *β*_1_ controls the steepness of the sigmoid function (inverse temperature) measuring the baseline sensitivity of choice to difference in decision variables, and *β*_0_ is the side bias with positive values corresponding to a bias toward choosing right. For the RL_Stim-Only_ model, *DV*_*Left*_ and *DV*_*Right*_ were assigned based on the stimulus identity appearing on the respective side for a given trial. For example, if stimulus A appeared on the left of the fixation, then *DV*_*Left*_ = *V*_*StimA*_ and *DV*_*Right*_ = *V*_*StimB*_. For the RL_Action-Only_ model, the decision variables (DV) simply corresponded to action values; i.e., *DV*_*Left*_ = *V*_*Left*_ and *DV*_*Right*_ = *V*_*Right*_.

#### Two-system models with stimulus- and action-based learning

As an extension of the above RL models, we considered hybrid RL models that constituted two value functions, *V*_Stim_ and *V*_Action_, to simultaneously track the reward value for alternative stimuli and actions and made choices based on a weighted sum of value signals from the two systems with a weight that was fixed on each block of the experiment (RL_Stim+Action_+Static *ω* or Static *ω* model for short). Specifically, the value functions were updated in parallel using Eqs. 3–4.

Therefore, the DV in this model was set to:

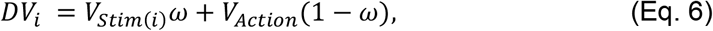

where *ω* represents the relative weight of the stimulus-based system compared to the action-based system, *i* ∈ {*Left, Right*}, and *V*_*Stim*(*i*)_ indicates the stimulus value for the option appearing on the side *i*. For example, if the stimulus A appeared on the left side, then *DV*_*Left*_ = *V*_*StimA*_ *ω* + *V*_*Left*_(1−*ω*) and *DV*_*Right*_ = *V*_*StimB*_ *ω* + *V*_*Right*_(1−*ω*). Similar to the learning rates and other parameters, a single value of *ω* was estimated for each block of trials.

Note that with this formulation of DV, the decision rule in Eq. 5 can be rewritten as (while dropping trial *t* for simplicity):

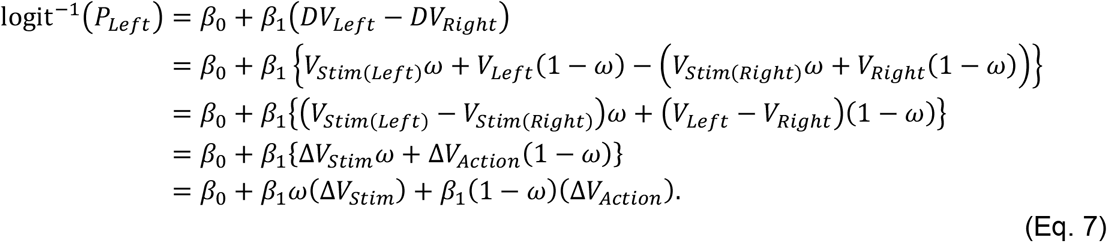

That is, *β*_1_*ω* and *β*_1_(1–*ω*) represent the sensitivity of choice to signals from the stimulus- and action-based systems, respectively. Therefore, *ω* controls the relative sensitivity of choice to the two competing systems with smaller *ω* corresponding to weaker influence of the stimulus-based system, and vice versa.

#### Two-system models with dynamic weighting

To allow for dynamic arbitration, we constructed hybrid models in which *ω* was updated on a trial-by-trial basis using the relative reliability of the two systems. In this model (Dynamic *ω*), the difference in reliability of the two systems is computed at the end of each trial to update the value of *ω* toward the more reliable system. More specifically, the relative reliability, *ΔRel*, between two systems at the end of trial *t* is computed as follows:

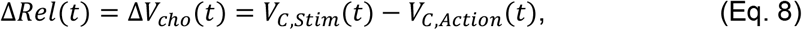

where *V*_C,Stim_ and *V*_C,Action_ correspond to the value of the chosen option in the stimulus- and action-based system, respectively, and *ΔV*_*cho*_ denotes the (signed) difference between the two. *ΔRel* ranges [-1, 1], with positive values indicating a more reliable stimulus-based system.

Intuitively, *ΔV*_*cho*_ signals the system that gives an overall larger value for the given choice and thus, is more reliable in predicting reward.

Subsequently, the relative sensitivity of choice to the two systems, *ω*, is updated as follows:

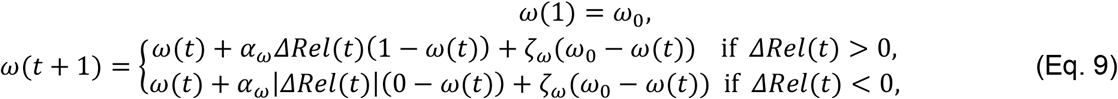

where *ω*_0_ is the initial *ω* on the first trial (onset of each block), *α*_*ω*_ is the baseline model arbitration rate, and ζ_*ω*_ is the passive decay rate that pulls *ω* toward its initial value *ω*_0_. This additional decay mechanism assumes that *ω* defaults back to its initial bias in the absence of exogenous input signaling the reliability difference (*ΔRel*). We focus on this model with passive decay for all analyses, as it fits better than the variants without the passive decay term (**Fig. S9A**). Importantly, the arbitration rates *α*_*ω*_ tend to be larger than the decay rates ζ_*ω*_ across groups and tasks (**Fig. S9B, C**).

#### Two-system models with dynamic weighting and separate baseline signals for stimulus- and action-based learning

To rule out the possibility that the observed effects in amygdala-lesioned monkeys are solely due to impairment in learning stimulus values (e.g., by reducing the strength of stimulus-value signals) and without any changes to arbitration processes, we included an additional parameter to separate these two types of changes. In the Dynamic *ω* model, an increase in the sensitivity to the stimulus-based system, *β*_1_*ω*(*t*), is strictly tied to a decrease in the sensitivity to the action-based system, *β*_1_(1–*ω*(*t*)), and vice versa. This constraint can be removed by introducing a constant factor which further scales the value of a given model before combining with the value from the other model. In this new model referred to as Dynamic *ω*-*ρ*, the decision variable is equal to:

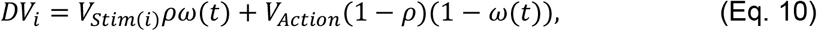

where *ρ* is a constant measuring the baseline ratio of signals from the stimulus-based to that from the action-based system independently of time-dependent arbitration weight (*ω*). The update for *ω* is the same as in the Dynamic *ω* model (Eq. 9). Therefore, similar to Eq.7, the decision rule for the Dynamic *ω*-*ρ* model can be simplified as follows:

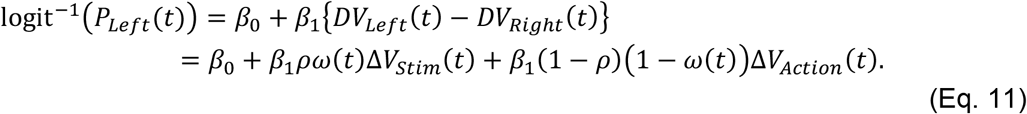

This shows that *β*_1_*ρ* and *β*_1_(1–*ρ*) can be seen as the baseline (maximum) sensitivity of choice to the stimulus- and action-based systems, respectively, before being modulated by the relative arbitration parameter *ω*. Therefore, including *ρ* allows for a variable amount of change (increase or decrease) to the sensitivity to the two learning systems, making the Dynamic *ω* model a special case of this model when *ρ* = 0.5. Here, *ρ* < 0.5 (*ρ* > 0.5) corresponds to low baseline activity or impairment in the stimulus-based (respectively, action-based) system. Because *ρ* is assumed to capture the baseline activity, we estimated a single value of *ρ* per each monkey for the entire duration of the experiment (see **Model fitting and cross validation** for more details). Because in this model, the arbitration weight is further weighted by *ρ* (or 1–*ρ*), we defined an “effective” arbitration weight to measure the overall relative weighting between the stimulus-based (*β*_Stim_= *β*_1_*ρω*) and action-based (*β*_Action_= *β*_1_(1–*ρ*)(1–*ω*)) systems as follows:

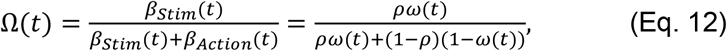

where *Ω* is the relative weight of the stimulus-based system with respect to the total sensitivity of the two systems. It is worth noting that the common baseline inverse temperature *β*_1_ is independent of *Ω*, which is determined only by *ρ* and *ω*, and that *Ω* reduces to *ω* when *ρ* = 0.5.

#### Alternative signals for estimating the reliability of learning systems

In addition to *V*_*chosen*_ as the reliability signal for updating *ω* in the dynamic models, we also considered several other quantities to estimate reliability. As the first alternative to *ΔV*_*cho*_ for the relative reliability used to update the relative weight (Eq. 8), we considered the difference in magnitudes of RPE (|*RPE*|) of the action- and stimulus-based systems:

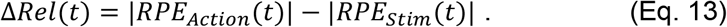

Conceptually, a system that yields better prediction of reward on a given trial has a lower magnitude of RPE and thus, is more reliable. Note that the difference in chosen stimulus and action values, *ΔV*_*cho*_, can be also rewritten as:

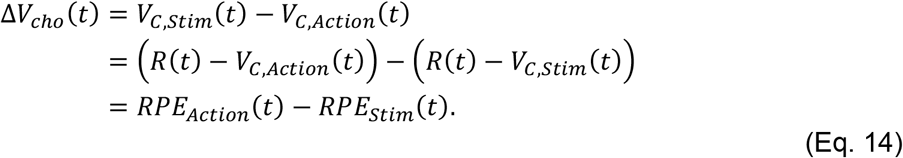

This demonstrates that using *ΔV*_*cho*_ for the relative reliability corresponds to the signed RPE instead of unsigned RPE.

We also considered the difference between the value of chosen and unchosen options within each system (|Δ*V*|) as a measure of the reliability of that system. Intuitively, this reliability signal is larger for a system that yields a better discernibility between the two competing options on a given trial. In this case, the relative reliability can be written as:

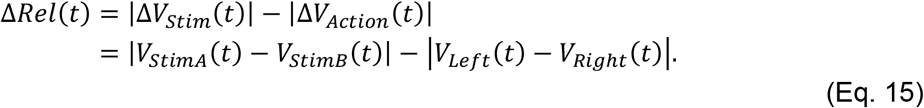

Finally, we also considered the total sum of the two value estimates (|Σ*V*|) as a signal for estimating the reliability of a given system. For this measure, reliability is larger for a system that gives overall larger combined values from the two options. In this case, the relative reliability is equal to:

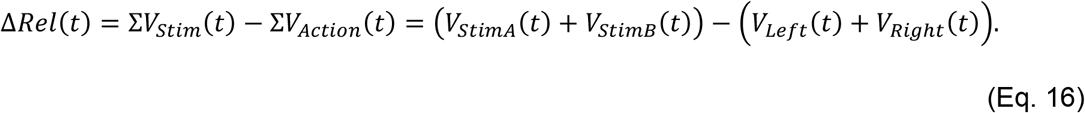

For all these different versions of *ΔRel* we used the same equation for updating *ω* (Eq. 9).

#### Effective arbitration rates

In the above formulation of arbitration mechanism, the rate of update for *ω* depends on several factors including the baseline *ρ*, baseline model arbitration rate *α*_*ω*_, the trial-by-trial difference in reliability (*ΔRel*), and the passive decay mechanism (ζ_*ω*_). In order to capture the overall transition rates in *Ω*, we computed the “effective” arbitration rates which quantify the effective rate of update given the direction of transition toward either the stimulus- or action-based system (**Fig. 3D**). More specifically, the effective arbitration rate on trial *t* when *Ω* increases, shifting choice toward the stimulus-based system, is defined as follows:

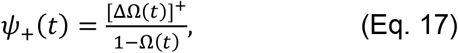

where *ψ*_+_ represents the effective arbitration rate toward the stimulus-based system on trial *t*, when *ΔΩ* > 0. Similarly, the effective transition rate when *Ω* decreases, biasing choice toward the action-based system, is defined as follows:

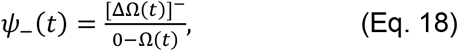

where *ψ*_-_ represents the effective arbitration rate toward the action-based system when *ΔΩ*< 0. *ΔΩ* = 0 corresponds to no change in arbitration.

### Model fitting, cross validation, and simulation

We used the standard maximum likelihood estimation method to fit choice data and estimate the best-fit parameters for the described models. One set of model parameters was fit to each session (consisting of ∼20 blocks) of monkeys’ choice data. Fitting was performed using the MATLAB optimization function *fmincon*, repeating the search for 100 initial random parameter values to ensure global minima.

For the Dynamic *ω*-*ρ* model, we assumed that the value of *ρ* is fixed for the entirety of the experiment and does not vary across sessions, as it aims to capture the relative strength of one learning system relative to the another. Accordingly, to estimate a single value of *ρ* for each monkey, we fitted the entire dataset of a given monkey and obtained a single set of best-fit parameters. From this set, we only kept the value of *ρ* and fit choice data again for the remaining parameters by allowing different values across sessions.

To compare the goodness of fit and determine the winning model, we used five-fold cross validation where each set of training/testing blocks was tested repeatedly with 50 unique instances. We created the training and testing sets as follows: for each subject, we categorized the blocks according to the permutation of each block type (*What-only, What, Where*) and reward schedule (80/20, 70/30, 60/40). Within each category, blocks were randomly assigned to training (80%) and testing (20%) sets. We then obtained best-fit parameters from the training set that minimized the negative log-likelihood across all training blocks and used these to calculate negative log-likelihood from each. We repeated this procedure 50 times for each category of blocks, each time using a unique combination of training and testing data. Final mean negative log-likelihoods were obtained by averaging across all instances within each block type.

To test whether the observed relationship between stimulus- and action-based learning indeed required competition between the two learning systems and was not due to task structure (**Fig. S4**), we simulated choice behavior using single-system or two-system models and computed correlations between simulated ERDS_Stim_ and ERDS_Action_. To that end, we used the fitted parameters to each session and simulated the choice behavior 100 times for each block, using the same random reversal position and reward schedule as the behavioral data. We obtained the final averaged ERDS values for stimulus and action.

### Models and parameters recovery

To perform model recovery (**Fig. S3A–D**), we simulated choice behavior of each model during a randomly created block environment similar to the experiment. Specifically, each session consisted of 30 blocks, each block having 80 trials with a reversal and with a randomly assigned reward schedule (80/20, 70/30, 60/40) and block type (What-only or What/Where). To ensure that choice behavior is simulated with a plausible range of parameters, we randomly sampled each parameter value from a kernel distribution fitted to all observed parameters values for a given model. We then fit all models and determined the best-fit model based on AIC for each simulated session. We repeated this procedure 1000 times and report the proportion of sessions that a given fitted model best accounts for each simulated model.

For parameter recovery (**Fig. S3E–F**), we simulated the choice behavior of the best model (Dynamic *ω*-*ρ*) using the estimated parameters from the experimental data, and then refit the simulated data with the same model. Each block environment was set up using the same reward schedule and reversal position as the actual experiment. We simulated each block one time to ensure that the total number of simulated trials is the same as that of the experiment, from which the true parameters were estimated. We recovered the parameters from the simulated data using the same fitting procedure, repeating the search for 100 initial random parameter values to ensure global minima. For recovering the *ρ* parameter, we used the entire simulated data for each monkey and obtained a single set of best-fit parameters as was done for the actual data.

### Data analysis and statistical tests

All analyses were carried out using MATLAB (MathWorks). All comparisons were performed using appropriate statistical tests reported throughout the text. For each test, we report the exact p-values and effect size, and Cohen’s *d* values when appropriate. For entropy metrics (ERDS) and estimated parameters of RL models, which are bounded between two values and often highly skewed, we used non-parametric statistical tests. All statistical tests used in this study were two-sided.

To categorize each trial as either *stimulus-* or *action-dominant*, we directly compared ERDS_Stim_ and ERDS_Action_ (computed from a moving window of ten trials). Trials with ERDS_Stim_ < ERDS_Action_ and ERDS_Action_ < ERDS_Stim_ were categorized as stimulus-dominant and action-dominant, respectively. We dropped trials with ERDS_Stim_ = ERDS_Action_, amounting to 11.1% of total trials in What-only and 11.8% in What/Where tasks. We used non-parametric tests (Wilcoxon rank sum test) to test whether two categories had significantly different RT as the distributions of RT were highly skewed.

To study the long-term adjustment in behavior across the time course of the experiment (**Fig. S6**), we calculated ERDS_Stim_, ERDS_Action_ for each block and regressed them on the proportion of the experiment completed (%) as the predictor variable within each subject. The results were then averaged across the subjects.

To estimate the effective arbitration rate across time (**Fig. 3E–G**), we computed the mean trajectory for each of two transition rates (toward stimulus- or action-based system) across blocks by aligning all trials relative to the beginning, reversal point, and the end of each block. For the bar plots in the insets, we computed the difference between the mean of the two transition rates within each block. In particular, we focused on the trials after the reversal to avoid potential confounds with initial bias (*ω*_*0*_) and observe the transition behavior after adjusting to the initial uncertainty of the block.

To plot performance (**Fig. 1E–G**) and effective arbitration rates over time (**Fig. 3E–G)**, we obtained the trajectories by concatenating 20 trials relative to the beginning, reversal point, and the end of each block, to account for the random reversal positions. These curves were then smoothed with a moving window of five trials, separately within the acquisition and reversal phase.

### Lead contact and materials availability

Further information and requests for resources should be directed to and will be fulfilled by the Lead Contact, Dr. Alireza Soltani. Analysis-specific codes are available at https://github.com/DartmouthCCNL/woo_etal_amygdala. Data are available upon request to the authors.

## Acknowledgments

We thank Chanc VanWinkle Orzell and Dmitriy Lisitsyn for their helpful comments on the manuscript. This work is supported by the National Institutes of Health (R01 DA047870 to A.S.) and by Intramural Research Program of the NIMH (ZIA MH002928).

## Author Contributions

J.H.W. and A.S. designed the study; V. C., C. T., K. R., and B. A. designed the experiments; J.H.W., V. C., C. T., K. R., B. A., and A.S. performed research; J.H.W. and A.S. analyzed data; J.H.W. and A.S wrote the first draft paper. All authors contributed to the revision of the paper.

## Supporting Information

### Supplementary Note 1

Considering previous findings on faster decision making during action-based compared with stimulus-based tasks^22^, we hypothesized that reaction time (RT) on a given trial depends on the learning system that controls the behavior more strongly on that trial. To test this hypothesis, we categorized trials as either stimulus- or action-dominant by directly comparing ERDS_Stim_ and ERDS_Action_ (computed from a moving window of ten trials). Trials that had ERDS_Stim_ < ERDS_Action_ were categorized as stimulus-dominant, and action-dominant if ERDS_Action_ < ERDS_Stim_.

For the What-only task in control monkeys, the vast majority of the analyzed trials (79.4%) was classified as stimulus-dominant (**Figure SN1A**). Interestingly, there was still some proportion of trials with ERDS_Action_ < ERDS_Stim_ (9.5%) even though action-based learning was irrelevant for performing the task. These action-dominant trials were marked by slightly slower RTs (M±SD = 227.7±91.8) than stimulus-dominant (220.9±73.5) trials (rank sum test; effect size *r* = -.0073, *p* = .0119). Comparing the performance for these two types of trials, we found that P(Better) was significantly lower for action-dominant (0.795) than stimulus-dominant (0.624) trials (two-sample z-test for proportions; *z* = 43.74, *p* < .001). These results suggest that most action-dominant trials happened when reward value estimates based on the two systems were close to each other, resulting in slower and more erroneous responses.

For the What/Where task, the proportions of stimulus-dominant and action-dominant trials were reflected in the respective block types: What blocks were marked by a higher proportion of stimulus-dominant trials (58.1%; **Figure SN1B**), whereas the majority of trials within Where blocks were categorized as action-dominant (61.5%; **Figure SN1C**). Critically, performance was higher for the correct strategy: in What blocks, P(Better) was significantly higher for stimulus-dominant (0.827) than action-dominant (0.59) trials (two-sample z-test for proportions; *z* = 118.7, *p* < .001), whereas in Where blocks, P(Better) was higher for action-dominant (0.798) than for stimulus-dominant (0.58) trials (*z* = -103.6, *p* < .001). In terms of RT, we found that for both block types, responses on stimulus-dominant trials were significantly slower than action-dominant trials (rank sum test; What: effect size *r* = .185, *p* < .001; Where: *r* = .166, *p* < .001). Consistently, categorization of blocks based on comparison of ERDS yielded a larger distinction in RTs than categorization simply based on block types, as reflected by a smaller effect size (rank sum test; What vs. Where: effect size *r* = .126, *p* < .001). These results show that entropy-based metrics could be used to identify the adopted model on a given trial and that RT reflected the adopted strategy, with a stimulus-based strategy recruiting consistently longer RT than an action-based one.

**Fig. SN1.**
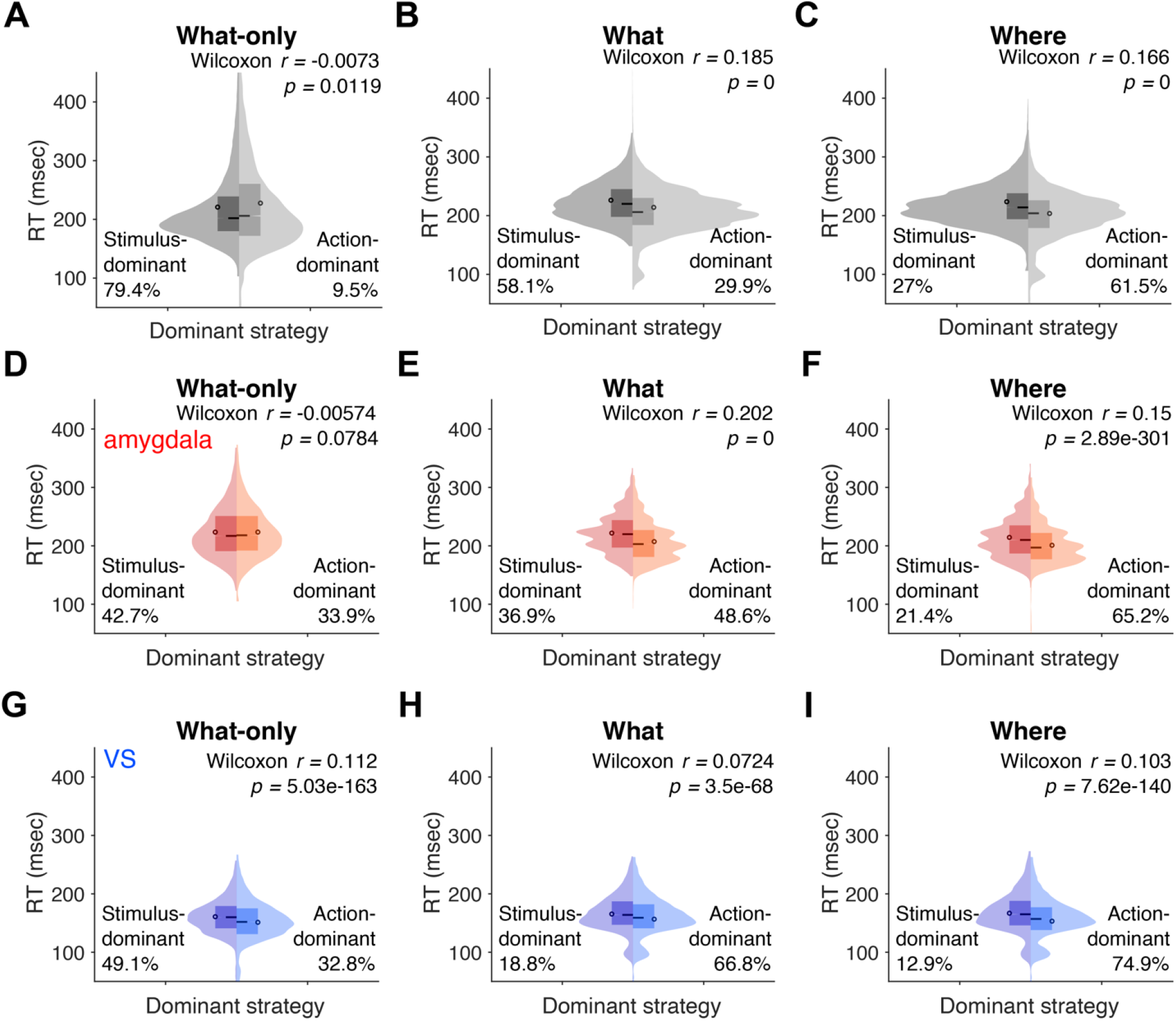
Distributions of reaction time (RT) for stimulus-dominant and action-dominant trials, separately for different tasks and block types. Each trial was categorized as either stimulus- or action-dominant by comparing ERDS_Stim_ and ERDS_Action_. Percentages indicate proportions of trials under each category (remaining percentages correspond to trials where both strategies were equally dominant). (**A– C**) RT data from control monkeys during What-only (A) and What/Where tasks (B,C). (**D–F**) RT data from amygdala-lesioned monkeys during What-only (D) and What/Where tasks (E,F). (**G–I**) RT data from VS-lesioned monkeys during What-only (G) and What/Where tasks (H,I). Circles in the violin plots represent means and black horizontal lines represent medians of the distributions. Reported are effect size *r* from Wilcoxon’s rank sum test and corresponding p-values. In the What/Where task (B–C,E–F,H–I), RT was significantly shorter for action-dominant trials.

**Fig. S1.**
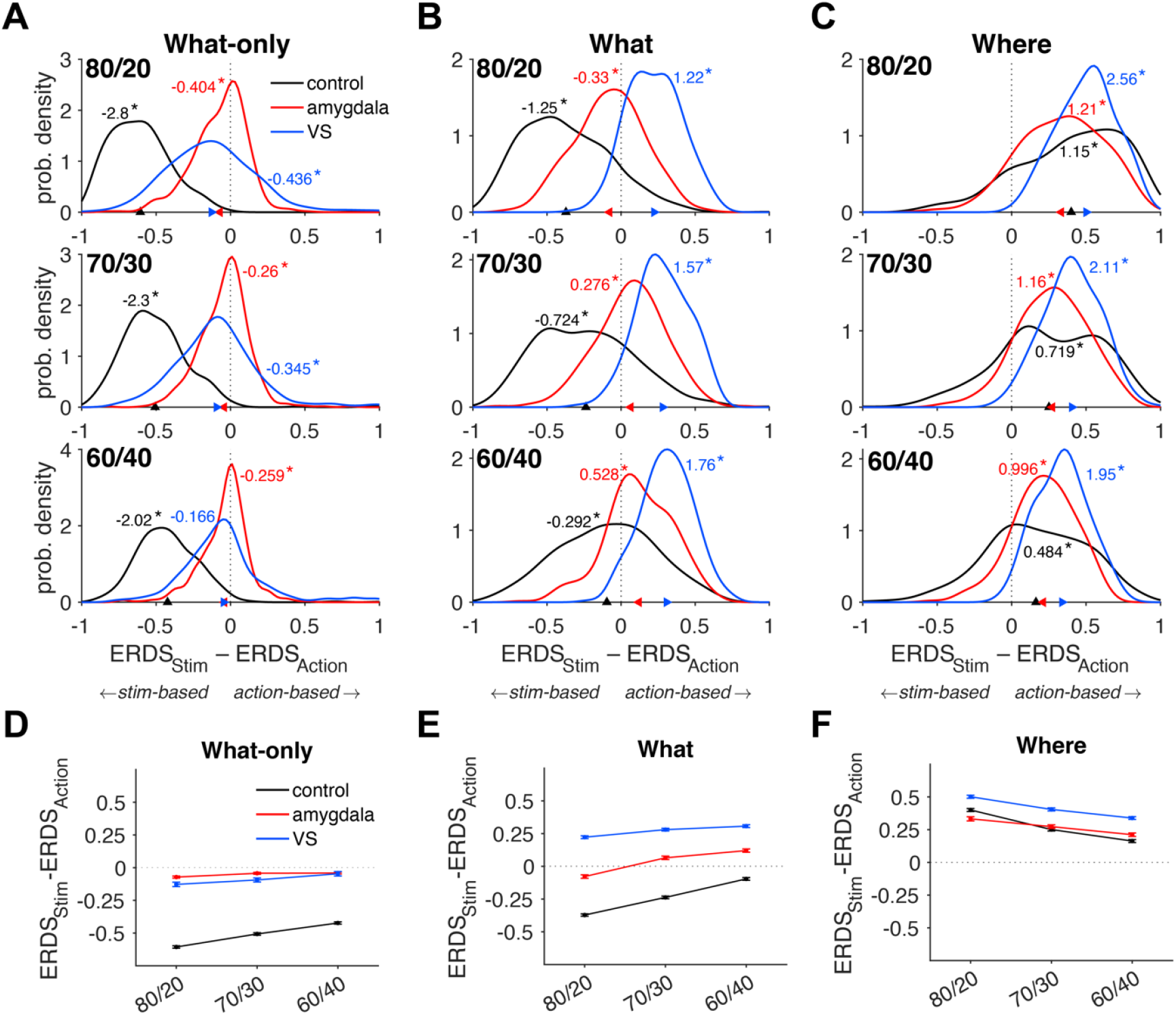
Interactions between stimulus-based and action-based learning depend on the uncertainty of the reward environment. (**A–C**) Distributions of the difference between ERDS based on stimulus identity and action (ERDS_Stim_ – ERDS_Action_) computed from the entire 80-trial blocks, separately for different reward schedules during What-only (A) and What/Where task (B, C). Reported are Cohen’s *d* values from paired-samples t-test between two metrics, and asterisks next to the value indicate significance (*p* < .001). Colors indicate control (black), amygdala-lesion (red), and VS-lesion (blue) monkeys. Triangles on the X-axis indicate means of the difference for each group. Negative ERDS_Stim_ – ERDS_Action_ values indicate dominance of the stimulus-based strategy, whereas positive values indicate dominance of the action-based strategy. In controls, means (black triangles) and Cohen’s *d* values become closer to zero as reward schedules are more uncertain. (**D–F**) Summary results for the panels in A–C. Plotted are mean values of ERDS_Stim_ – ERDS_Action_, by each group and reward schedule during What-only (D) and What/Where (E,F) tasks. Larger reward uncertainty leads to increasing influence of the competing (incorrect) learning strategy in a given block type as revealed by ANOVA: What-only controls: *F*(2,1666) = 63.2, *p* = 3.61×10^−27^; What-only amygdala: *F*(2,1531) = 0.19, *p* = .829; What-only VS: *F*(2,896) = 3.3, *p* = .0372; What controls: *F*(2,3004) = 190.3, *p* = 1.55×10^−78^; What amygdala: *F*(2,1531) = 63.9, *p* = 8.50×10^−27^; What VS: *F*(2,896) = 16.7, *p* = 7.96×10^−8^; Where controls: *F*(2,2934) = 118.1, *p* = 4.70×10^−50^; Where amygdala: *F*(2,878) = 18.0, *p* = 2.24×10^−8^; Where VS: *F*(2,896) = 53.0, *p* = 2.09×10^−22^. Only amygdala-lesioned monkeys during the What-only task did not show this effect. Overall effect sizes (Cohen’s *d*) across all reward schedules were as follows: What-only controls = -2.24, What-only amygdala = -0.309, What-only VS = -0.32, What controls = -0.699, What amygdala = 0.15, What VS = 1.48, Where controls = 0.758, Where amygdala = 1.1, Where VS = 2.09.

**Fig. S2.**
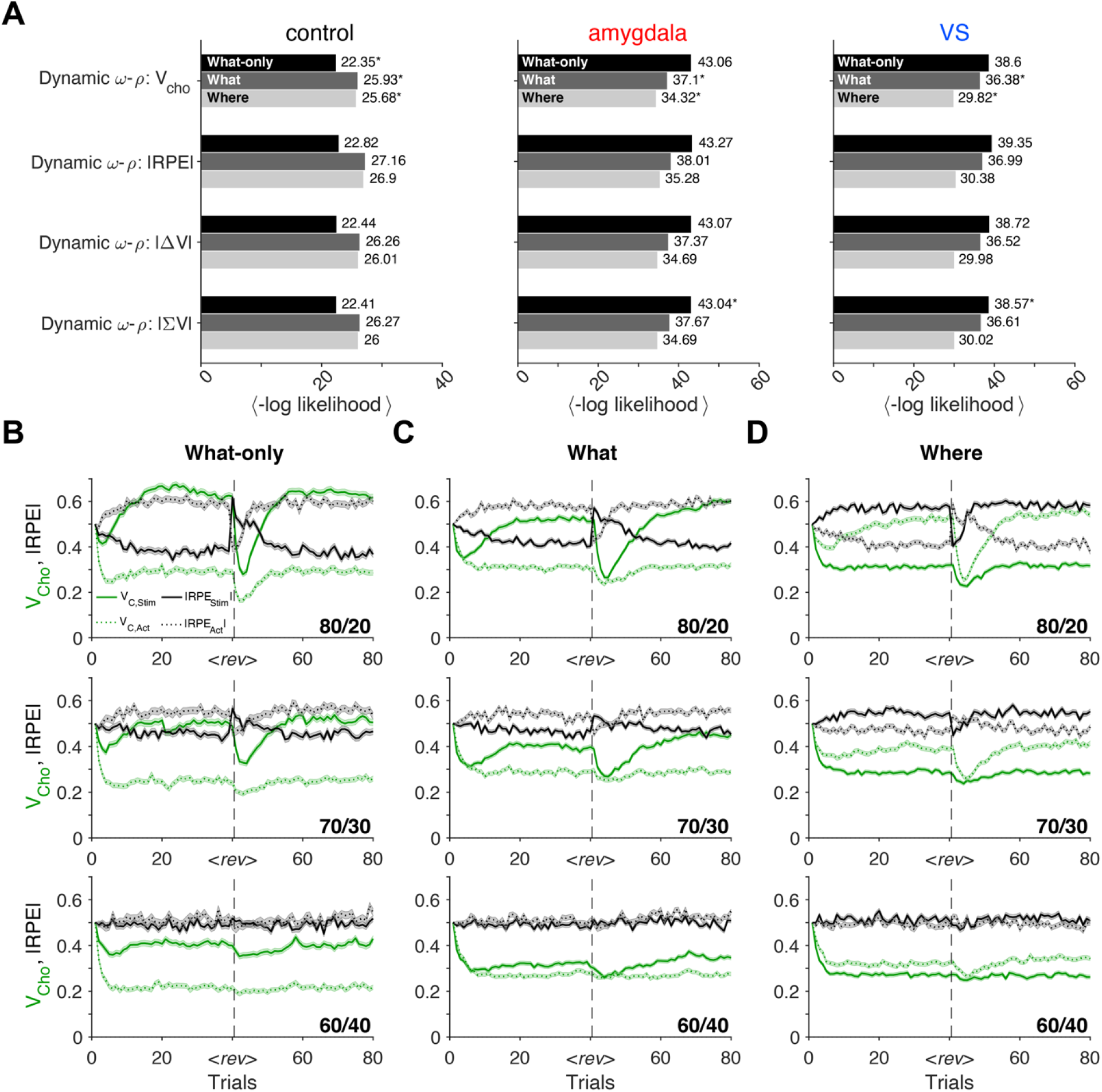
Comparison of different reliability signals for dynamic arbitration between the stimulus- and action-based systems. (**A**) Goodness of fit using five-fold cross-validation of the Dynamic *ω*-*ρ* model with different types of reliability signals. V_cho_: value of chosen option V_cho_, |RPE|: magnitude of RPE, |ΔV|: magnitude of difference between chosen and unchosen options, ΣV: sum of chosen and unchosen options. Reliability signal based on V_cho_ consistently yields the best fit to choice behavior across all block types in control animals. (**B–D**) V_cho_ signal provides better separability between reliable and unreliable systems. Plotted are control monkeys’ averaged trajectories for the value of the chosen option (V_cho_, green) and magnitude of RPE (|RPE|, black) within each system (stimulus-based for solid, and action-based for dotted lines) as a function of trial number during What-only (B), What (C), and Where (D) blocks. Each row corresponds to each reward schedule: Top = 80/20, middle = 70/30, bottom = 60/40. In these plots, we used estimated parameters from the Static *ω* model (RL_Stim+Action_ +Static *ω*) to avoid confounds in signals in dynamic models.

**Fig. S3.**
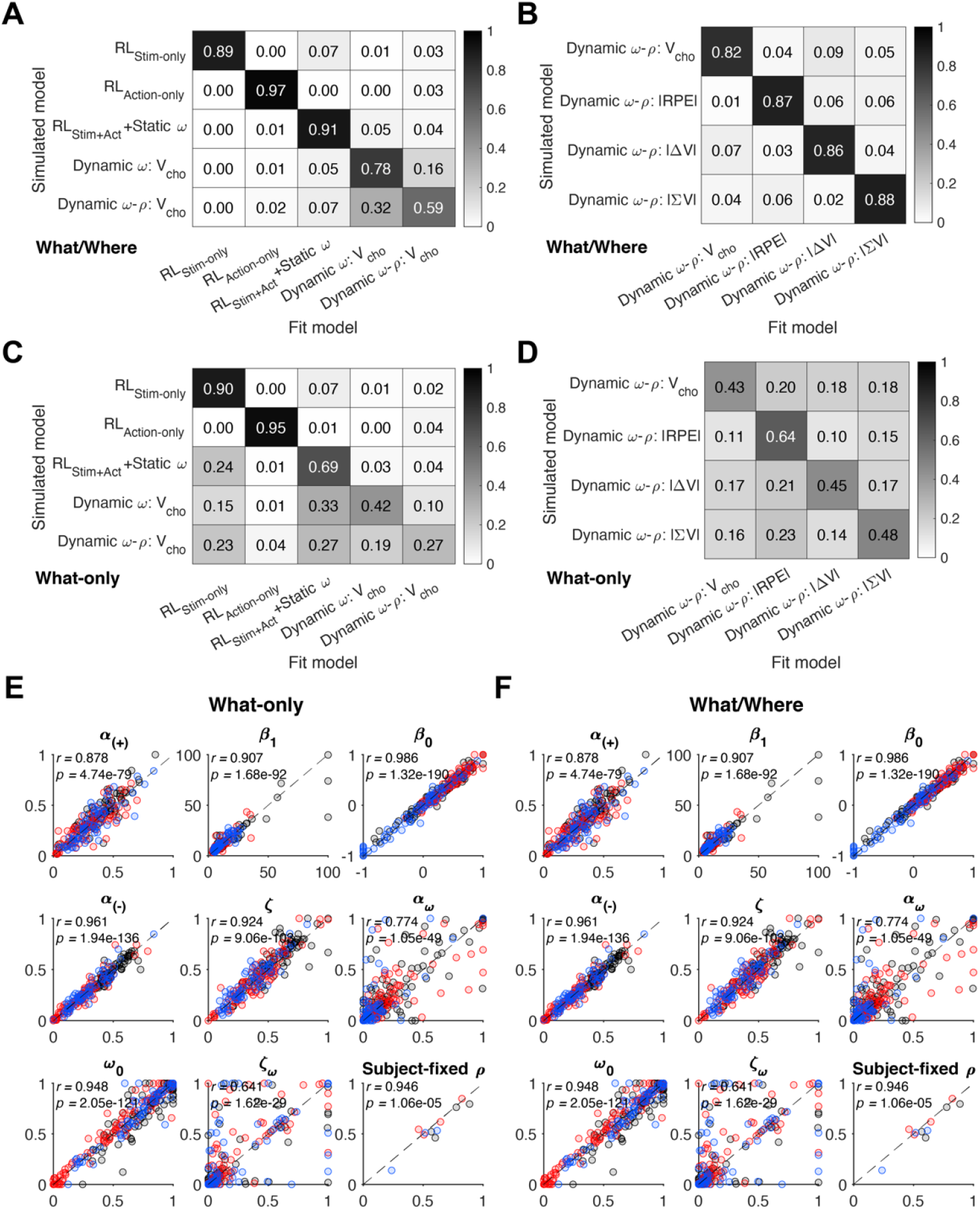
Models and parameters recovery. (**A**) Model recovery of one- and two-system models, with and without dynamic arbitration. Reported numbers are proportion of sessions that are best explained by a given fitted model (minimum AIC) for each simulated model. The Dynamic *ω*-*ρ* model is less distinguished from the Dynamic *ω* model, as the latter is a special case of the former model with *ρ* = 0.5. (**B**) Model recovery for alternative reliability signals in dynamic arbitration, as shown in **Figure S2**. (**C–D**) Same plot as in A–B but for What-only task. In this task, simulated behaviors of more complex models become less distinct from their simpler counterparts. (**E–F**) Scatter plots of true (X-axis) and recovered (Y-axis) parameters from the Dynamic *ω*-*ρ* model. Each panel shows each parameter of the model for the three groups of monkeys (controls: black; amygdala-lesioned: red; VS-lesioned: blue). Reported are Pearson’s correlation between true and recovered parameters. All correlations were *r* > 0.60 and significant (*p* < .001). See **Figures S8** and **S9** for detailed description of the fitted parameters. *ρ* is assumed to be fixed for each monkey, whereas all other parameters are estimated for each session.

**Fig. S4.**
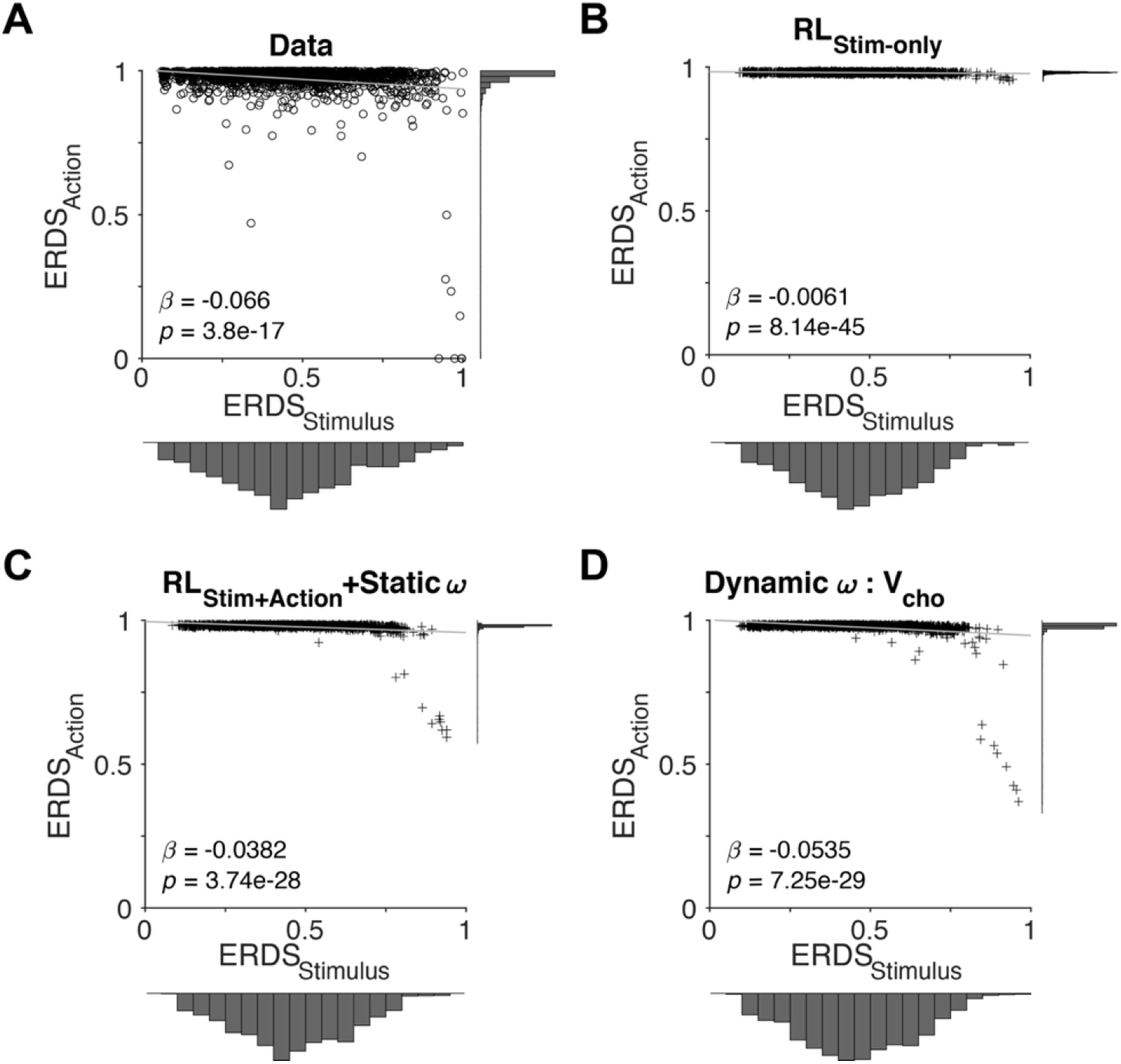
Comparison of empirical and simulated ERDS in control monkeys during the What-only task provides evidence for presence of multiple learning systems. (**A**) Scatter plot of entropy of reward-dependent strategy based on stimulus identity (ERDS_Stim_, X-axis) vs. action (ERDS_Action_,Y-axis) in control monkeys during the What-only task. Reported are regression coefficient (*β*) and its p-value for regression of ERDS_Action_ on ERDS_Stim_. Histograms in the X- and Y-axis show distributions of ERDS_Stim_ and ERDS_Action_, respectively. (**B**) Simulated metrics using estimated parameters of the model with stimulus-learning system only (RL_Stim-Only_). (**C**) Simulated metrics using estimated parameters of the static two-system model (RL_Stim+Action_+Static *ω*). (**D**) Simulated metrics using estimated parameters of the two-system model with dynamic adjustment using V_chosen_ as the reliability measure (Dynamic *ω*: V_cho_). The Dynamic *ω* model better captures the variability in ERDS_Action_ compared to the Static *ω* model shown in panel C.

**Fig. S5.**
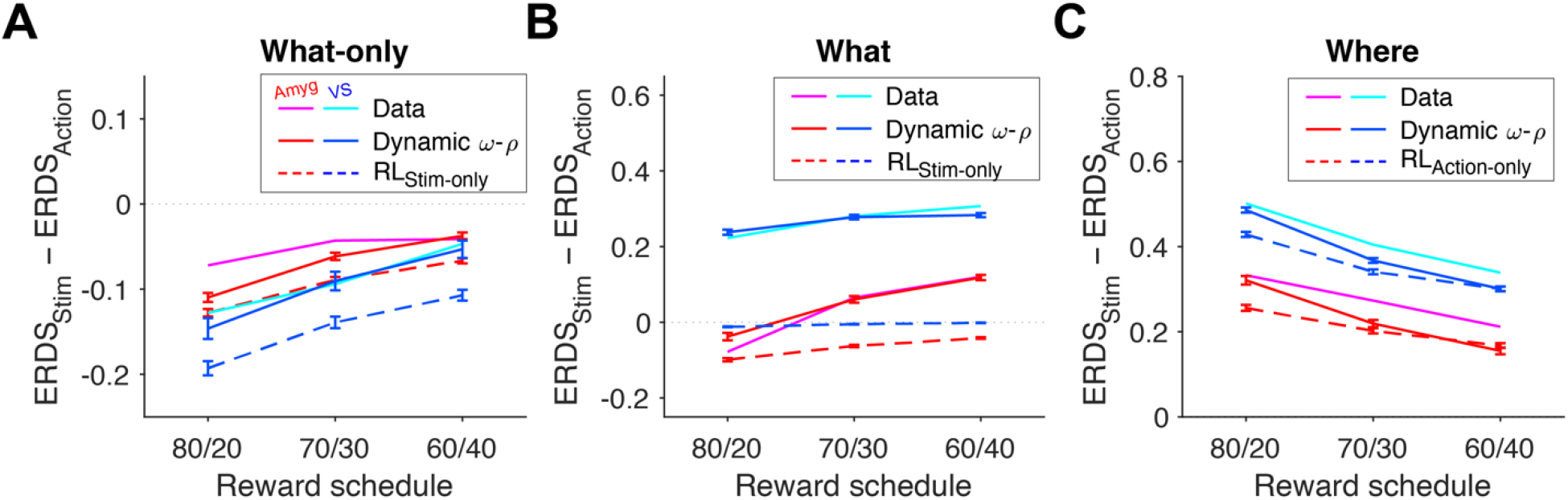
Model validation results demonstrating that the best model captures the relative strength of the two strategies better than the model with no arbitration. (**A**) Plotted are mean values of ERDS_Stim_ – ERDS_Action_ from the empirical data in lesioned monkeys (same as those shown in **Figure S1D**, magenta and cyan lines) and model simulation (solid and dashed lines). The two-system model with dynamic arbitration, the Dynamic *ω*-*ρ* model, more effectively captures the data compared to the model without arbitration (RL_Stim-Only_). Error bars = s.e.m. Error bars for data are not shown to avoid clutter. (**B**) Empirical and simulated values of ERDS_Stim_ – ERDS_Action_ for What blocks during the What/Where task. Conventions are the same as in A. (**C**) Same plot as in B but for Where blocks during the What/Where task. Choice behavior for one-system model is simulated with the RL_Action-Only_ model.

**Fig. S6.**
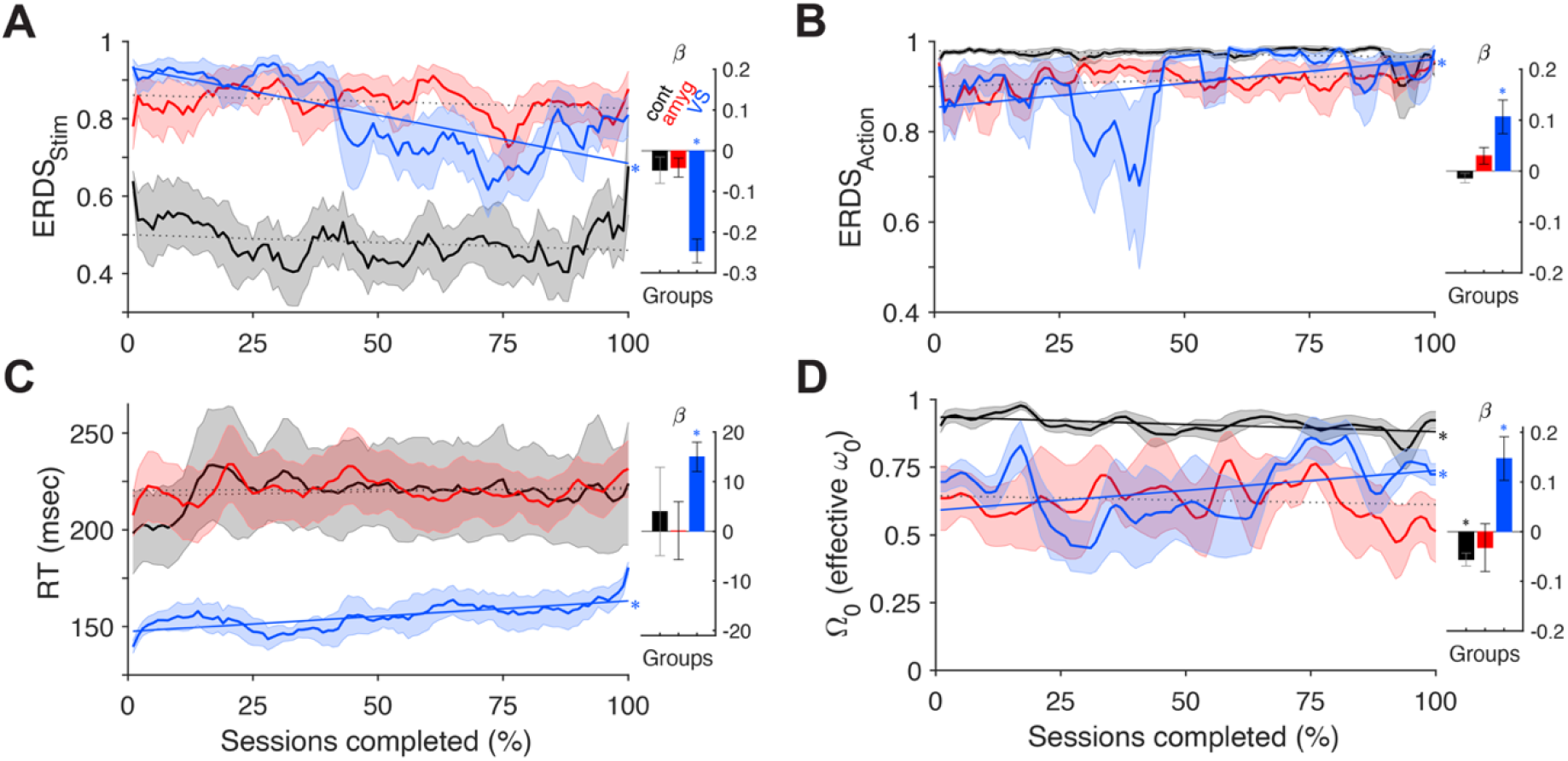
Contribution of the amygdala to behavioral adjustments over long timescales. (**A–B**) Time course of entropy of reward-dependent strategy on stimulus identity (ERDS_Stim_, A) and performed action (ERDS_Action_, B) for controls (black), amygdala- (red), and VS-lesioned (blue) monkeys during the What-only task. Number of blocks completed was normalized by each monkey into percentages (error bar = SEM across subjects). Straight lines indicate least-squares lines regressing ERDS on the fraction (%) of sessions completed. Asterisks indicate significance of regression (*p* < .001). Sub-panel to the right shows the bar plot of the slope of the fitted line for each group. (**C–D**) Time course of median RT (C) and initial effective arbitration weight *Ω*_0_ (D) within each block for the entirety of the What-Only task. Conventions are the same as in panels A–B.

**Fig. S7.**
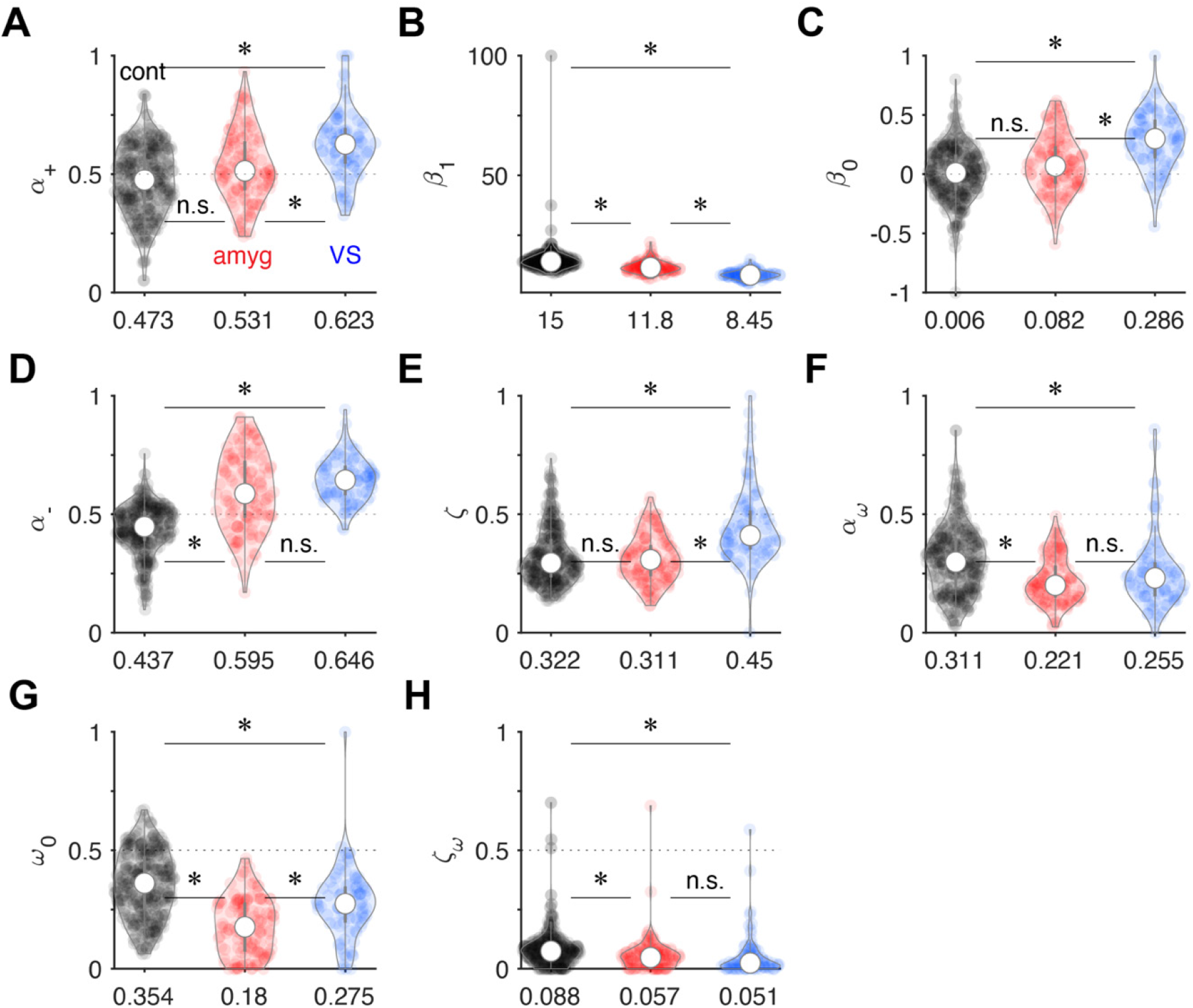
Distributions of estimated model parameters in What/Where task across three animal groups. Plotted are the distributions of estimated parameters of the best model (Dynamic *ω*-*ρ* with V_cho_) fitted to the choice behaviors of controls (black), amygdala- (red), and VS-lesioned (blue) monkeys during the What/Where task. *α*_+_: learning rate on rewarded trials (A). *β*_1_: common inverse temperature for stimulus- and action-based systems (B). *β*_0_ : side bias term, positive if preferring right option (C). *α*_-_: learning rate on unrewarded trials (D) ζ: decay or forgetting rate for the unchosen option (E). *α*_*ω*_: arbitration transition rate (F). *ω*_0_: initial arbitration weight on the first trial of each block (G). ζ_*ω*_: decay rate for arbitration weight *ω* toward initial value (H). Asterisks indicate significant difference between groups by Wilcoxon rank sum test (*p* < .001; n.s. = not significant). Circles in the violin plots indicate medians, and the numbers on the X-axis indicate mean values of the distributions. *ρ* parameter is assumed to be fixed for each subject and is shown in **Figure 4A**.

**Fig. S8.**
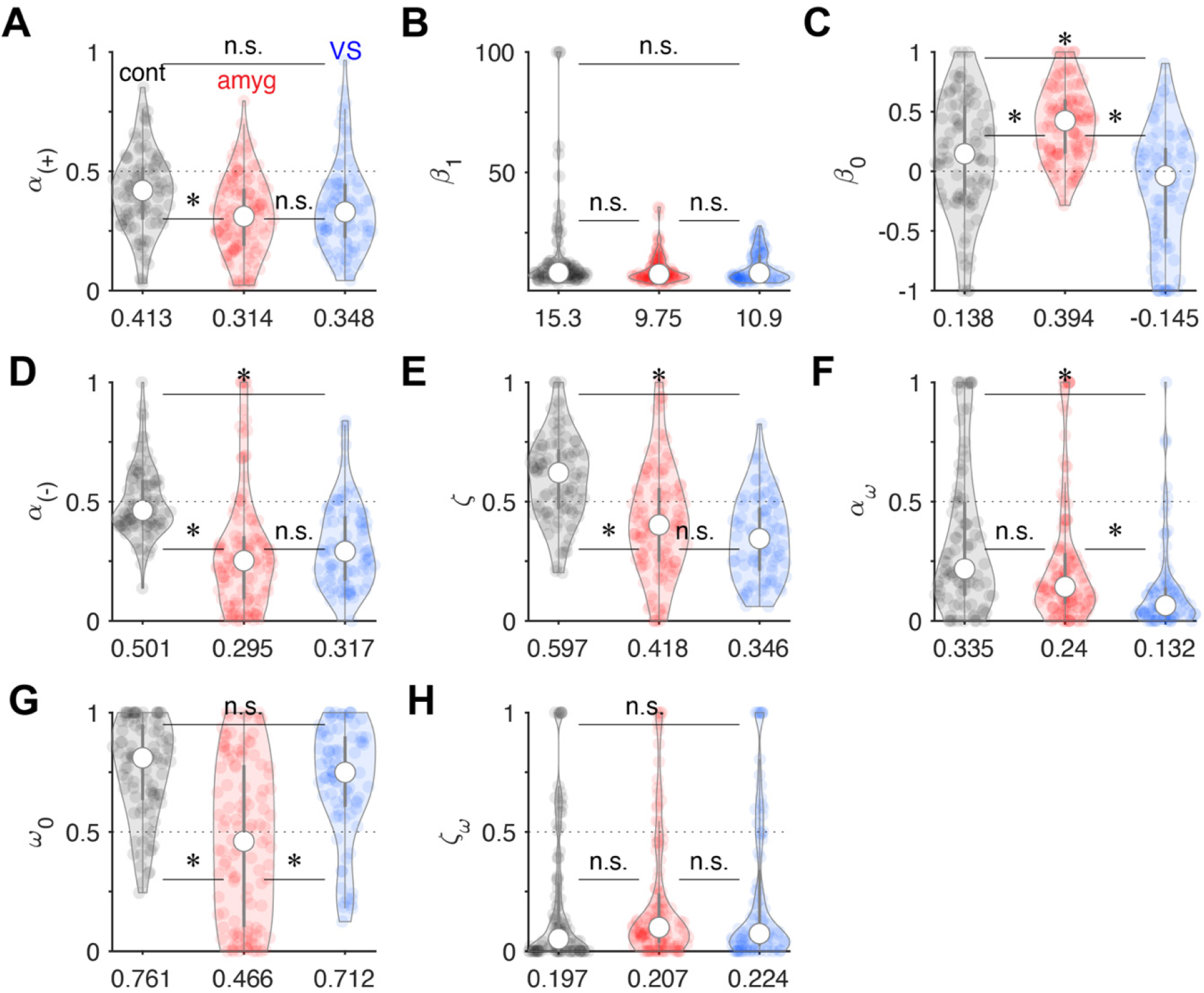
Distributions of estimated model parameters in What-only task across three animal groups. Plotted are the distributions of estimated parameters of the best model (Dynamic *ω*-*ρ* with V_cho_) fitted to the choice behaviors of controls (black), amygdala-(red), and VS-lesioned (blue) monkeys during the What-only task. *α*_+_: learning rate on rewarded trials (A). *β*_1_: common inverse temperature for stimulus- and action-based systems (B). *β*_0_: side bias term, positive if preferring right option (C). *α*_-_: learning rate on unrewarded trials (D) ζ: decay or forgetting rate for the unchosen option (E). *α*_*ω*_: arbitration transition rate (F). *ω*_0_: initial arbitration weight on the first trial of each block (G). ζ_*ω*_: decay rate for arbitration weight *ω* toward initial value (H). Asterisks indicate significant difference between groups by Wilcoxon rank sum test (*p* < .001; n.s. = not significant). Circles in the violin plots indicate medians, and the numbers on the X-axis indicate mean values of the distributions. *ρ* parameter is assumed to be fixed for each subject and is shown in **Figure 4A**.

**Fig. S9.**
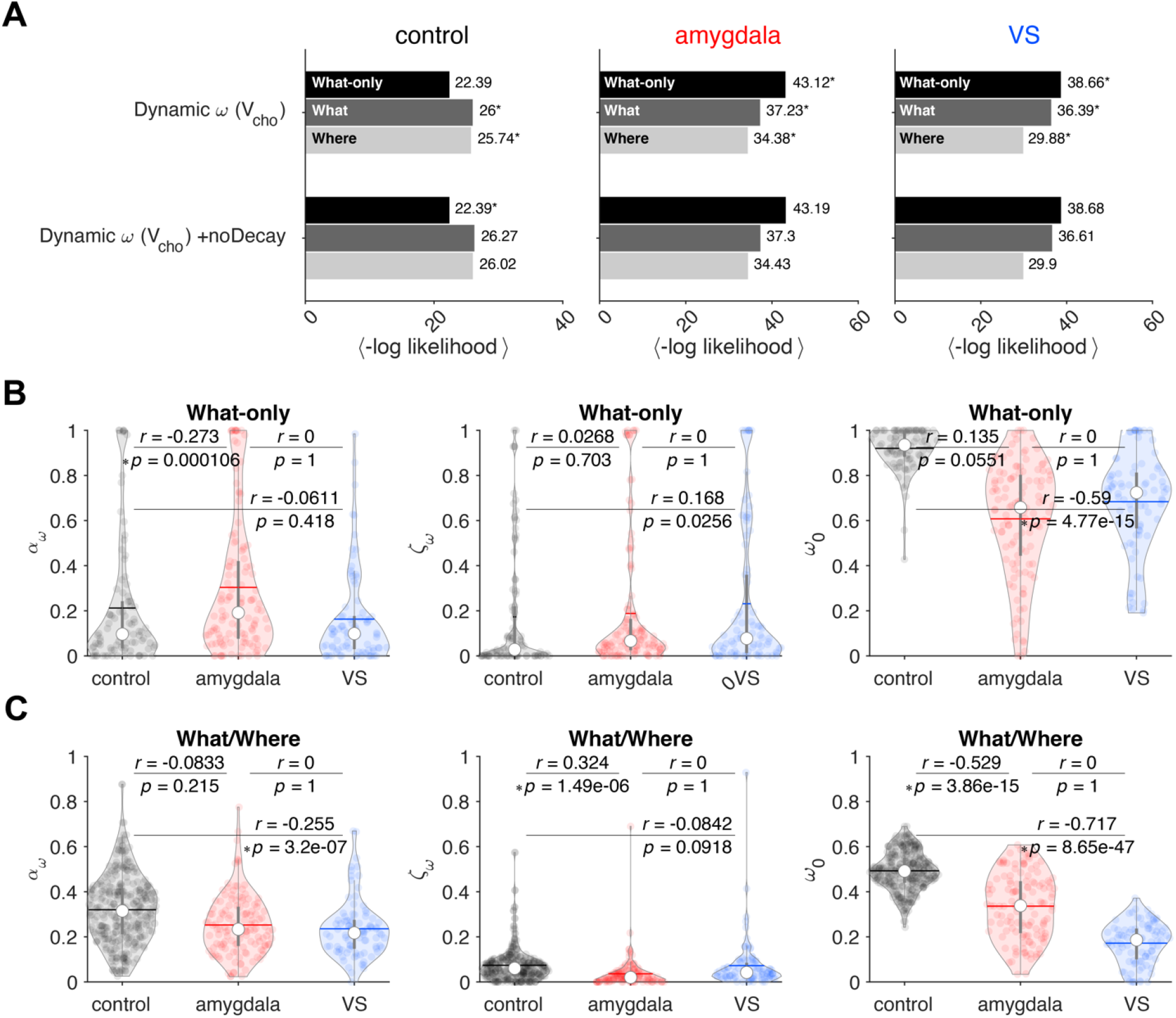
Comparison between dynamic models with or without passive decay in arbitration weight, and estimated parameters in the better-fitting model. (**A**) Goodness of fit using five-fold cross-validation of dynamic models with or without passive decay for *ω*. Model with the additional passive decay mechanism better accounts for the choice behavior in all groups, especially during the What/Where task. (**B**) Distribution of estimated parameters for sessions in controls (black), amygdala (red), and VS (blue) groups during the What-only task, showing parameters for *α*_*ω*_ (model arbitration/update rate, left), ζ_*ω*_ (decay rate for *ω*, middle), and *ω*_0_ (initial bias or baseline decay value for *ω*, right). Asterisks show significance from Wilcoxon rank sum tests between groups, with effect size *r* and its p-value. Circles = median, horizontal lines = means. Arbitration rates *α*_*ω*_ tended to be larger than the passive decay rate ζ_*ω*_ across groups (Wilcoxon signed-rank test; controls: *z* = 2.15, *p* = .032, effect size *r* = 0.158; amygdala: *z* = 4.92, *p* = 8.73×10^−7^, effect size *r* = 0.32; VS: *z* = -0.192, *p* = .848, effect size *r* = -0.015). (**C**) Same plot as in (B) but for the What/Where task. Arbitration rates *α*_*ω*_ were significantly larger than the passive decay rate ζ_*ω*_ across all groups, suggesting that *α*_*ω*_ is the primary source of the transitions in *ω* (Wilcoxon signed-rank test; controls: *z* = 15.03, *p* = 4.74×10^−51^, effect size *r* = 0.603; amygdala: *z* = 9.91, *p* = 3.84×10^−23^, effect size *r* = 0.612; VS: *z* = 7.82, *p* = 5.45×10^−15^, effect size *r* = 0.583).

## Notes

### Competing Interest Statement

The authors have declared no competing interest.

